# Nanomechanics of wild-type and mutant dimers of the tip-link protein protocadherin 15

**DOI:** 10.1101/2023.10.17.562769

**Authors:** Camila M. Villasante, Xinyue Deng, Joel E. Cohen, A. J. Hudspeth

## Abstract

Mechanical force controls the opening and closing of mechanosensitive ion channels atop the hair bundles of the inner ear. The filamentous tip link connecting transduction channels to the tallest neighboring stereocilium modulates the force transmitted to the channels and thus changes their probability of opening. Each tip link comprises four molecules: a dimer of protocadherin 15 and a dimer of cadherin 23, all of which are stabilized by Ca^2+^ binding. Using a high-speed optical trap to examine dimeric PCDH15, we find that the protein’s configuration is sensitive to Ca^2+^ and that the molecule exhibits limited unfolding at a physiological Ca^2+^ concentration. PCDH15 can therefore modulate its stiffness without undergoing large unfolding events in physiological Ca^2+^ conditions. The experimentally determined stiffness of PCDH15 accords with published values for the stiffness of the gating spring, the mechanical element that controls the opening of mechanotransduction channels. When PCDH15 has a point mutation, V507D, associated with non-syndromic hearing loss, unfolding events occur more frequently under tension and refolding events occur less often than in the wild-type protein. Our results suggest that the maintenance of appropriate tension in the gating spring is critical to the appropriate transmission of force to transduction channels, and hence to hearing.

## Introduction

The transformation of sound waves into neural signals, a process known as mechanoelectrical transduction, occurs in the cochlea of the internal ear. To allow humans to hear with the range and precision that we do, this process must be both sensitive and adaptive. The apical surface of each hair cell bears actin-filled stereocilia arranged in order of height. Each stereocilium is connected by a filamentous tip link to its tallest neighbor^1^. When the hair bundle is deflected towards its tall edge by a sound stimulus, the stereocilia pivot and tense the gating springs that control the opening of the mechanoelectrical transduction channels atop each stereocilium. When sufficient tension has been conveyed, the channels open and allow cations within the endolymphatic fluid surrounding the hair bundles to flow into the cells and transduction to occur^2^ (Figure 1A).

**Figure 1.**
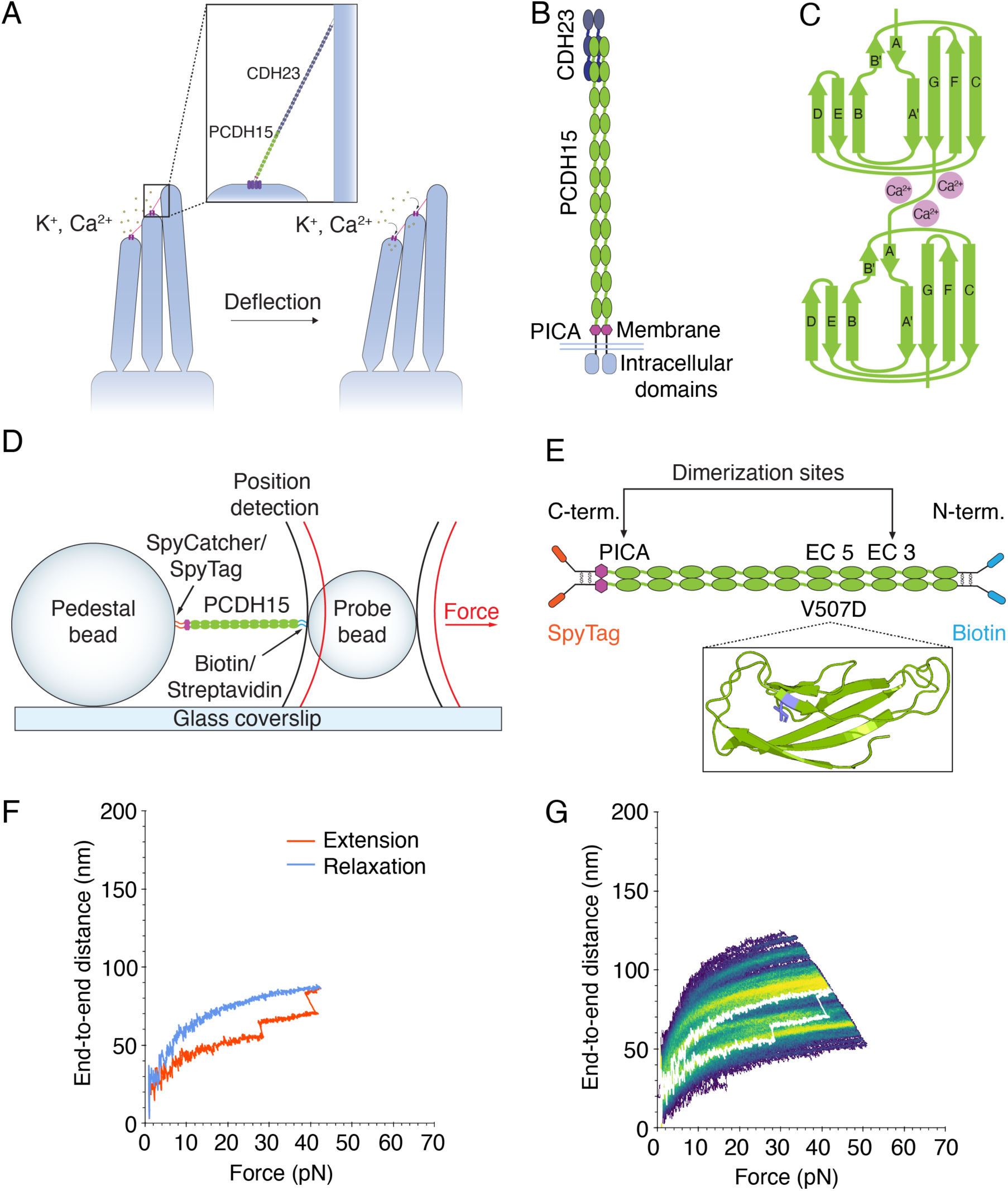
The structure of PCDH15 and measurements with an optical trap. (A) When stereocilia are deflected towards the tall edge of a hair bundle, the tip links (inset) connecting them stretch, opening the mechanotransduction channels (purple) atop each stereocilium and allowing the ions within the endolymph to flow into the cell, resulting in depolarization. (B) PCDH15 comprises 11 EC domains and a PICA domain at its carboxy terminus. PCDH15 binds with CDH23 at its amino terminus in a handshake interaction. (C) EC domains are composed primarily of β-sheets. Ca^2+^ binding in the linker regions and at the edges of the domains stabilizes the structure against unfolding. (D) In our apparatus, the protein is tethered between two beads and two laser beams act on the probe bead to measure its position and exert force on it. (E) In our experimental construct, we maintained the dimerization sites in PCDH15 while adding two disulfide bonds to each end in addition to the distinct molecular tags at each end. The V507D construct was identical to the wild-type construct save for the insertion of the mutation in place of V507 (purple in inset). (F) A force-ramp experiment comprises the extension phase of the cycle, during which force is increased at a constant rate, and the relaxation phase of the cycle, during which force is decreased back to a minimum at the same constant rate. In these experiments, the minimum force is 1 pN and 2 s elapse between successive cycles. Unfolding events can be seen as sudden steps during an individual extension. (G) Repetition of a force-ramp cycle hundreds of times on the same protein molecule yields a heat map in which the brighter colors represent more highly occupied states. The heatmap superimposes both extension and relaxation phases of every cycle. The illustrative cycle from panel F is overlaid in white.

It is plausible that the gating springs are the filamentous tip links that connect the transduction channels of a stereocilium to its tallest neighbor. Each tip link comprises four protein molecules^3^: a dimer of protocadherin 15 (PCDH15) and a dimer of cadherin 23 (CDH23). Because abolishing the tip links halts mechanotransduction^4^ whereas allowing the tip links to regenerate restores sensitivity^5^, the tip link—or the attachments at its two ends—likely controls the opening and closing of the channels and serves as the gating spring. Moreover, the hundreds of mutations in tip-link proteins that result in human hearing loss^6–10^ underscore the role of the tip link in hearing.

The direct association of PCDH15 with the transduction-channel complex by way of its transmembrane and cytoplasmic domains^11^ implicates the protein in channel gating. PCDH15 comprises 11 extracellular cadherin (EC) domains and a protocadherin 15-interacting, channel-associated (PICA) domain, also known as MAD12^12^ or EL^11^ (Figure 1B). The EC domains, which are composed of Greek key motifs, are similar in their folding patterns (Figure 1C). The intervening linker regions can bind up to three Ca^2+^ ions^12^ and stabilize the entire molecule against unfolding by force^13^. The concentration of Ca^2+^ in the cochlear endolymph is approximately 20 μM, a value much lower than that in the rest of the body^14^. The dissociation constant of Ca^2+^ at the linker regions lies in the range of tens to hundreds of micromolar^12^, a range that might confer additional modulation of PCDH15 behavior at physiological Ca^2+^ concentrations.

Depending on the local characteristic frequency and whether measured in inner or outer hair cells, the enthalpic stiffness of gating springs in the rat’s cochlea^15^ lies between 0.5 mN·m^-1^ and 4 mN·m^-1^. Although monomers of PCDH15 have an enthalpic stiffness of approximately 10 mN·m^-1^, the measured stiffness of monomeric PCDH15 is lower over the physiological range of forces^16^. This softening represents the contribution of entropic elasticity, which stems from the extension of the interdomain linker regions, the EC domains themselves, and any unfolded portions of the molecules. We asked here whether dimeric PCDH15 would exhibit a similar softening over the physiological range of forces and whether its enthalpic stiffness would accord with that of the gating spring. To understand whether PCDH15 has the appropriate properties to be a component of the gating spring, we measured its stiffness, inquired what factors control its mechanical responses, and examined how it softens under force.

## Results

### Experimental conditions

To probe the mechanical properties of PCDH15 directly, we used an optical trap with sub-nanometer spatial resolution and microsecond temporal resolution^16,17^ (Figure 1D). In a typical experiment, PCDH15 was tethered between two beads: at its carboxy terminal it was attached by a SpyTag-SpyCatcher bond^18^ to a pedestal bead covalently attached to a glass coverslip, and at its amino terminal it was linked thorough a biotin-streptavidin interaction to a probe bead in solution. The anchors that attached PCDH15 to the beads on either end were separated from the protein by short, unstructured peptides. Two lasers acted on the probe bead: a highly stable 1064 nm position-sensing laser, which detected the three-dimensional position of the probe bead and thus the extension of the construct, and an 852 nm force-producing laser, which exerted force on the probe bead and therefore on the protein.

To explore the range of physiological forces that PCDH15 experiences in the ear^19^, we conducted force-ramp experiments in which force was increased at a constant rate from a resting level of 1 pN and then decreased at the same rate to a resting level of 1 pN, where it was held for 2 s before the next cycle. During these extension-relaxation cycles, the protein sometimes underwent conformational changes. Such unfolding events could be seen as steps in the end-to-end distance (Figure 1F). After a force-ramp cycle had been repeated up to hundreds of times on a single protein molecule, all the cycles could be displayed as a heatmap in which lighter colors correspond to more frequently occupied states and darker colors represent trajectories that occurred only once or infrequently (Figure 1G).

Because of the importance of Ca^2+^ to the structure of PCDH15, we performed our experiments at three representative Ca^2+^ concentrations: 3 mM, a saturating level meant to populate all the Ca^2+^-binding sites; 20 μM, a physiological concentration in the mammalian cochlea^20–22^; and with no Ca^2+^, but in the presence of the Ca^2+^ chelator EDTA. Previous studies on monomeric PCDH15 showed that its mechanical behavior changes with the Ca^2+^ concentration^16^, so we explored whether the dimer shows a similar Ca^2+^ dependence.

### Ca^2+^ sensitivity of PCDH15 under physiological forces

PCDH15 naturally dimerizes at EC3 and the PICA domain^23^. To ensure that force was distributed equally to both strands of each PCDH15 dimer, we devised a construct in which the monomers are attached to one another at each end by paired disulfide bonds derived from the Fc hinge region of human immunoglobulin^24^ (Figure 1E). This study used murine PCDH15, which has high sequence homology^25^ to human PCDH15.

As exhibited both by a representative cycle (Figure 2A) and by the single bright branch on a representative heatmap (Figure 2B), dimeric PCDH15 underwent little unfolding at a saturating level of Ca^2+^. At low forces, PCDH15 extended easily in response to applied force, a behavior that reflects entropic elasticity owing to the straightening of the molecule’s thermal undulations. At higher forces, after most entropic degrees of freedom had been pulled out, the relationship between the end-to-end distance and the applied force became nearly linear. The remaining extensibility resulted from the enthalpic or Hookean stiffness of PCDH15.

**Figure 2.**
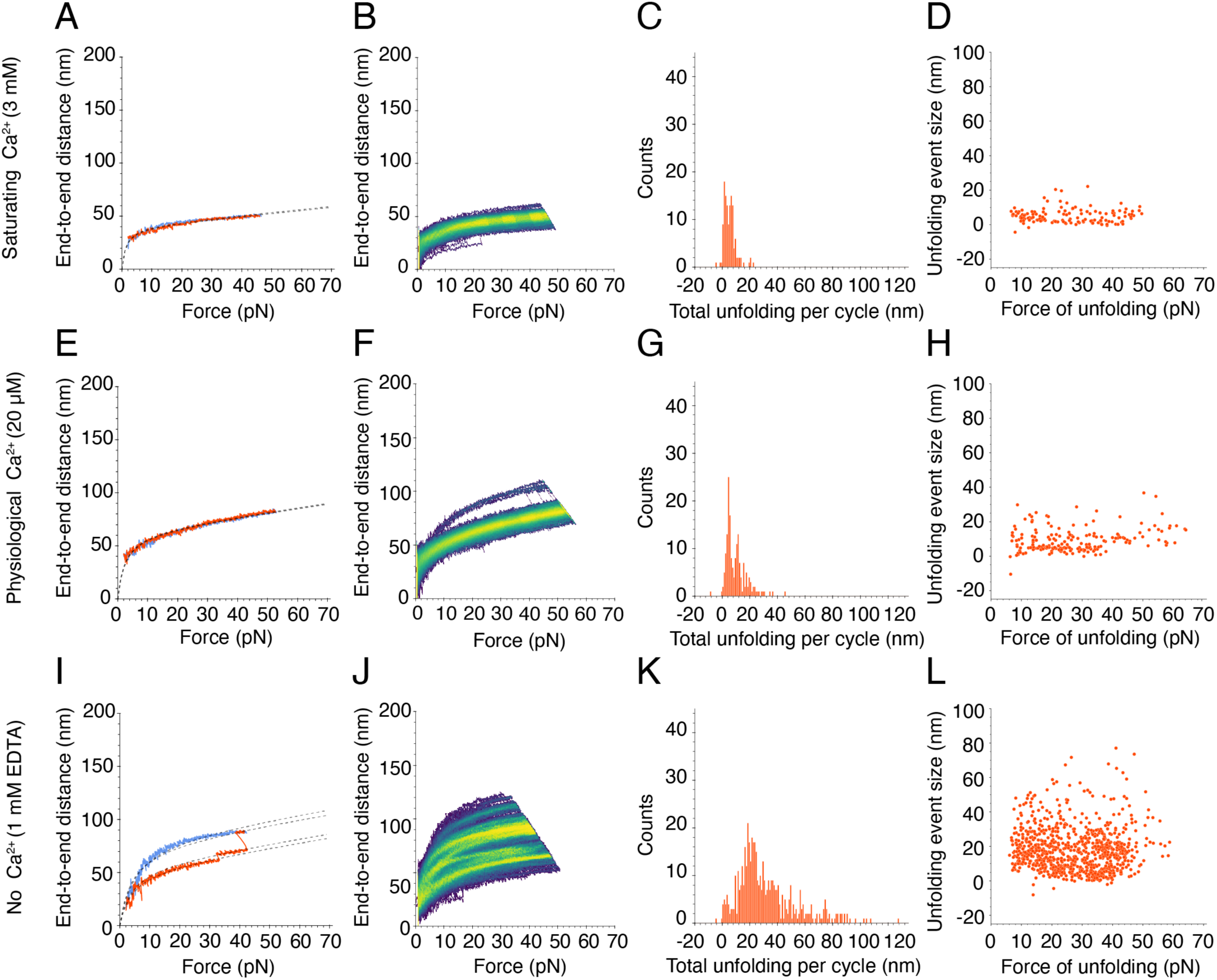
Force-ramp responses of wild-type PCDH15 at three Ca^2+^ concentrations. (A) At a saturating Ca^2+^ concentration of 3 mM, PCDH15 unfolds infrequently. The dashed line represents a fit of our model, Equation (1), to the extension and relaxation phases of the illustrative cycle. (B) In a heatmap for a saturating level of Ca^2+^, 3 mM, the single bright branch indicates one highly occupied state that reflects the infrequent and small unfolding events. (C) The total unfolding length during the extension phases of all cycles in all datasets at 3 mM Ca^2+^ peaked at 1.8 ± 0.1 nm and 6.3 ± 0.2 nm (means ± SEMs; *N =* 5 datasets; *n =* 131 events). (D) Many unfolding events occurred at forces below 10 pN at 3 mM Ca^2+^. (E) At a physiological Ca^2+^ concentration of 20 μM, PCDH15 often does not unfold and is more extensible than under 3 mM Ca^2+^ in the same force range. (F) The illustrative heatmap has one bright branch indicative of the infrequent and small unfolding events at 20 μM Ca^2+^. (G) At 20 μM Ca^2+^, the total unfolding per cycle was slightly greater than at a saturating level of Ca^2+^, with peaks at 4.5 ± 0.1 nm and 11.5 ± 0.1 nm (means ± SEMs; *N =* 4 datasets; *n =* 172 events). (H) At 20 μM Ca^2+^, individual unfolding events with a mean of 4.6 ± 0.1 nm occurred predominantly at forces below 40 pN. (I) An illustrative force-ramp cycle in the absence of Ca^2+^ and in the presence of 1 mM EDTA shows a small unfolding event followed by a larger one. In this case, our protein model, Equation (1), is fitted separately to each segment demarcated by the unfolding events. (J) The numerous bright branches in the heatmap reflect the multiple preferred conformational states of PCDH15 in the absence of Ca^2+^ and in the presence of 1 mM EDTA. (K) The total unfolding per cycle peaked at 23.3 ± 0.6 nm (mean ± SEM; *N =* 6 datasets, *n =* 490 events), but many larger events occurred in the absence of Ca^2+^ and in the presence of 1 mM EDTA. (L) There is no clear relationship between the size of individual unfolding events and the corresponding forces in the absence of Ca^2+^ and in the presence of 1 mM EDTA.

To quantify and classify the unfolding changes that occurred, we used a saturation model augmented with a linear-spring term: if *x* is the end-to-end distance of the PCDH15 construct,

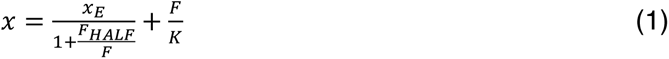

The maximal entropic extension of PCDH15 is given by *x*_E_; *F*_HALF_ is the force at which entropic extension is halfway complete. The contribution of enthalpic stiffness is given by the second term, the extension of a linear spring of stiffness *K* under force *F*. The value of *K* was determined by averaging the inverse spatial derivatives at forces exceeding 30 pN for every cycle of all data sets at each Ca^2+^ concentration.

At a saturating concentration of Ca^2+^, 3 mM, few discrete unfolding events occurred, and their magnitudes were relatively small. The frequency distribution of the size of these unfolding events was bimodal with peaks at 2.0 nm and 6.6 nm (Figure S6). We asked whether these individual unfolding events happened in succession within the same cycle, which would suggest the occurrence of a larger unfolding event through a multi-step process. We summed the total unfolding lengths in the extension phase of each cycle, during which the majority of unfolding events occurred. The total unfolding length per cycle was predominantly below 20 nm, with a bimodal frequency distribution peaking at 1.8 nm and 6.3 nm (Figure 2C). These events could correspond to unfolding of the inter-domain linker regions, which—because they lack secondary structure—are likely to be the first components of PCDH15 to unfold under force. Individual linker regions would give rise to an additional length between 1 nm and 2 nm when extended, so it is plausible that the unfolding of linker regions yielded the unfolding events that we observed. Furthermore, not all the linker regions bind three ions of Ca^2+^—some bind none whereas others bind one or two ions^12^—so we might have observed extension of these linker regions even at a saturating level of Ca^2+^. It is also possible that the small extensions reflected the straightening of the kinked EC9-10 linker^12,26^, which does not bind Ca^2+^ and could extend up to 4 nm. We also asked whether there was any relationship between the size of individual unfolding events and the force at which they occurred. We found that unfolding events around the 2.0 nm and 6.6 nm frequency peaks occurred more often at forces between 10 pN and 20 pN than at forces below and above this interval (Figure 2D).

PCDH15 also unfolded infrequently at a physiological Ca^2+^ concentration of 20 μM. Nevertheless, at that concentration PCDH15 was strikingly more extensible than at a saturating level of Ca^2+^ (Figure 2E,F). In other words, despite the scarcity of unfolding events, the dimer was softer at the lower concentration of Ca^2+^. This finding was surprising, for we had expected an increase in compliance to emerge through more frequent unfolding events. This observation might reflect the differential Ca^2+^-binding behaviors of the linker regions in PCDH15: the heterogeneity in the linkers, combined with the relatively low physiological concentration of Ca^2+^ and the binding affinity for Ca^2+^, might result in the overall softening. As was observed at a saturating Ca^2+^ concentration, the frequency distribution of total unfolding length per cycle was bimodal with peaks at 4.5 nm and 11.5 nm (Figure 2G). These values were similar in magnitude to the peaks of total unfolding length per cycle that we observed at a saturating level of Ca^2+^. It was not possible to discern the origin of these classes of unfolding events owing to the repetitive structure of PCDH15. However, from the magnitudes of the conformational changes and assuming a length of 0.40 nm per amino acid^27^, we conjecture that these two classes arose from the inter-domain linker regions of PCDH15 that, when unfolded, would have given rise to 16 nm of additional length. The individual unfolding events were again distributed across a range of forces (Figure 2H). These results suggest that, at saturating and physiological Ca^2+^ conditions, unfolding of the linkers confers flexibility to PCDH15 and modulates its stiffness.

In the absence of Ca^2+^, PCDH15 underwent many types of unfolding that can be seen both in individual cycles and in numerous highly occupied branches on the heatmap (Figure 2I,J). There were far more unfolding events than were observed at higher Ca^2+^ concentrations. The sum of the lengths of unfolding events per cycle, although peaking in frequency at 23.3 nm (Figure 2K), reached values above 100 nm. We would expect the unfolding of a full-length cadherin domain to extend the end-to-end length of PCDH15 by 33 nm to 45 nm, depending on the particular domain^28^ that unfolded, and assuming a length of 0.40 nm per amino acid^29^ less 4.5 nm to account for the loss of the folded domain^30^. Because we do not observe a clear peak in that range, we infer that the unfolding of full domains was not the primary response to applied force. The length of unfolding at peak frequency that we observed could correspond to partial domain unfolding: if the A and B strands of an individual EC domain were to come undone, the end-to-end distance would increase by approximately 17 nm. The individual unfolding events occurred over a range of forces, with no clear relationship between the size of the unfolding event and the force at which it occurred (Figure 2L). This result is surprising, because we had expected to see more frequent domain unfolding in the absence of Ca^2+^, but our results suggest that an alternative mechanism is in play.

### Effects of a hearing-loss mutation on PCDH15

Numerous mutations in the tip link result in hearing loss^6,7,10,31^. Such mutations can cause either syndromic deafness, in which deafness is accompanied by other deficits such as blindness or vestibular dysfunction, or non-syndromic deafness, which involves exclusively hearing loss. Though PCDH15 is present in the retina and vestibular labyrinth as well as the cochlea, individuals with non-syndromic deafness retain normal retinal and vestibular function: only their hearing is affected, which raises interesting questions about the pathophysiology of these mutations.

We sought to understand how a deafness-causing mutation affects the mechanics of PCDH15. We chose to study V507D, the murine homolog of the human V528D variant that is associated in humans with non-syndromic, autosomal, recessive deafness type 23 (DFNB23)^31^. This point mutation of a highly conserved valine in EC5 was identified in a Newfoundland family whose members exhibit prelingual hearing loss. The V507 residue occurs in the β-sheet of the B strand of EC5 (Figure 1E), which is not a site of dimerization, nor part of the handshake interaction with CDH23, nor part of the Ca^2+^-binding sites, any of which would result in obvious disruptions of tip link integrity. We then predicted the structures of wild-type and V507D EC5 domains using AlphaFold2 Colab^32–36^. Alignment of the crystal structure and AlphaFold2- predicted wild-type EC5 showed that the prediction was highly accurate (Figure S4A). The predicted structure of V507D EC5 indicates that the β-sheet structure of the B strand would be disrupted by the substituted negatively charged aspartic acid residue (Figure S4B,C). This alteration could result in an easier unfolding of strands A and B, which in the native protein are stabilized by a parallel β-sheet interaction between the A and G strands and a stronger anti- parallel β-sheet interaction between the B and E strands (Figure 1C). Disruption of the β-sheet interaction of strands B and E could therefore cause an instability of strands A and B. We conjecture that the change of this hydrophobic valine to a negatively charged aspartic acid weakens the force-bearing ability of the β-sheet and thus compromises the structural integrity of EC5. Furthermore, the EC5-6 linker region binds only one Ca^2+^ ion, which might predispose EC5 to unfolding.

As the concentration of Ca^2+^ decreased from saturating to absent, PCDH15 V507D exhibited a striking pattern of increasing dysregulation. At a high Ca^2+^ concentration of 3 mM, and in contradistinction to the wild-type dimer, there were two populations of V507D molecules. Some V507D molecules appeared to behave more like wild-type PCDH15 at the same saturating level of Ca^2+^, with the minimal unfolding observed in individual cycles reflected by the single bright branch on the heatmap (Figure S5A,B). Other V507D molecules exhibited more unfolding during individual cycles, revealed by multiple bright branches on the heatmap (Figure 3A,B). We included both kinds of force-extension trajectories in further analysis, for the tethering statistics (see Supplementary Information) gave us confidence that individual trajectories corresponded to individual molecules rather than multiple tethers, and we observed both kinds of trajectories often enough to deem them significant.

**Figure 3.**
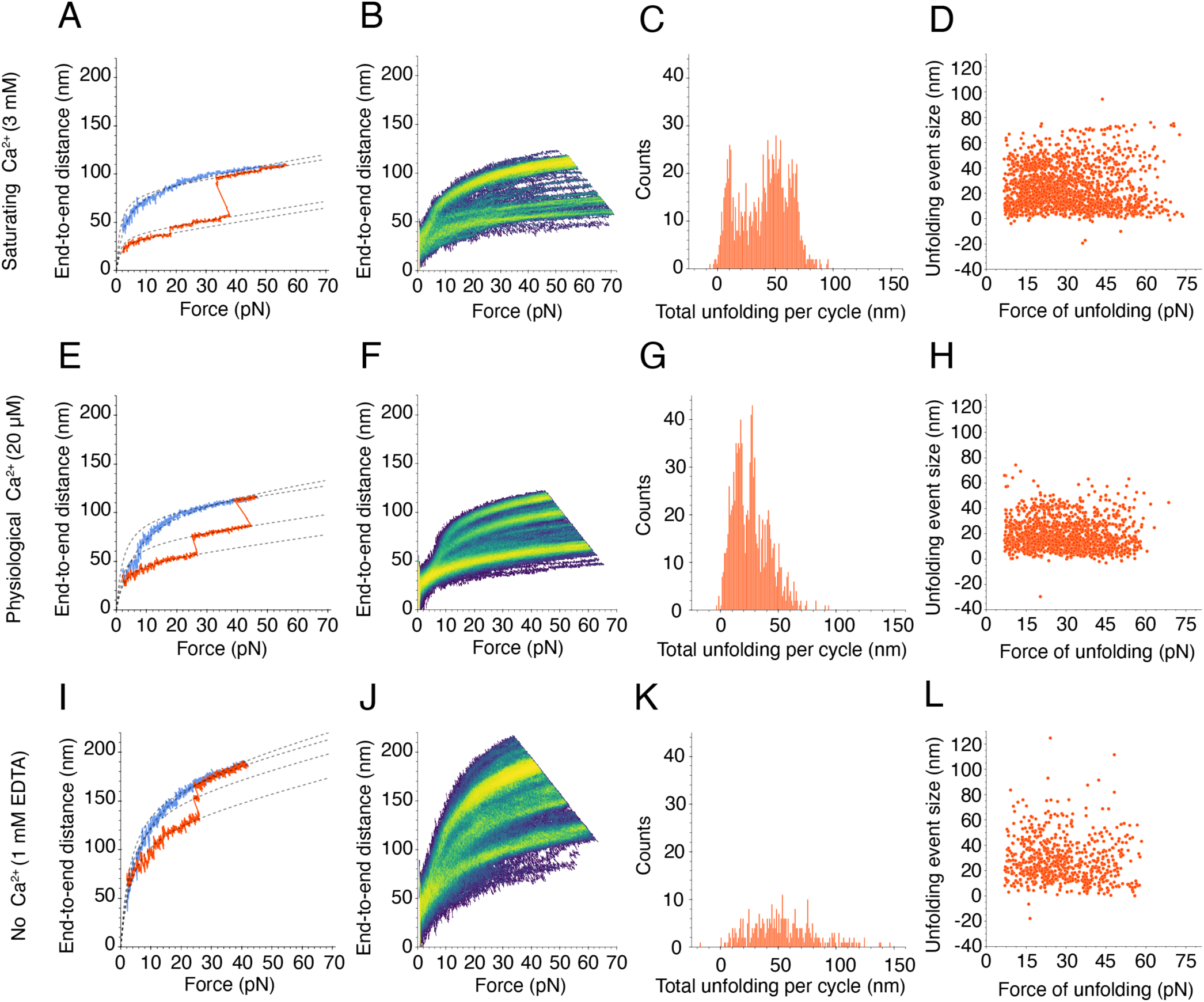
Force-ramp responses of V507D at three Ca^2+^ concentrations. (A) At a saturating level of Ca^2+^, 3 mM, a subset of V507D molecules underwent only small unfolding events while another subset had more frequent unfolding (Figure S5A). The dashed line represents the fit of our model, Equation (1), to the segmented extension and relaxation phases. (B) The bright branches on the heatmap reflect the frequent unfolding seen in the individual cycle. (C) The frequency distribution of total unfolding length per cycle peaked around 8.8 ± 0.1 nm, 50.0 ± 0.2 nm, and 66.3 ± 0.1 nm (means ± SEMs; *N* = 24 datasets; *n* = 1889 events). (D) As the size of the individual unfolding event got larger, the force of unfolding tended to be lower, with many events occurring around 30 pN. (E) At a physiological concentration of Ca^2+^, 20 μM, frequent unfolding was seen on the single cycle level. (F) The unfolding seen in the individual cycles is reflected by the bright branches on the exemplary heatmap. (G) The frequency distribution of total unfolding length per cycle during the extension phase peaked around 14.2 ± 0.2 nm, 27.3 ± 0.1 nm, and 39.4 ± 0.3 nm (means ± SEMs; *N* = 16 datasets; *n* = 1081 events). (H) As at a saturating level of Ca^2+^, the larger unfolding events were associated with smaller forces of unfolding. (I) When Ca^2+^ was absent, V507D extended easily, even without large unfolding events. (J) The bright branches on the illustrative heatmap reflect the unfolding behavior seen on the individual cycle level. The upper limits of the end-to-end distance at this level of Ca^2+^ exceeded those seen at higher concentrations of Ca^2+^. (K) The frequency distribution of total unfolding length per cycle was unimodal, with one peak at 49.3 ± 1.4 nm (mean ± SEM; *N* = 4 datasets; *n* = 362 events). (L) In the absence of Ca^2+^, there was no clear relationship between the size of individual unfolding events and the forces at which they occurred.

Considering both populations, we observed many more unfolding events per cycle from V507D than from wild-type PCDH15 at the same saturating concentration of Ca^2+^, for which we largely observed only small unfolding events below 10 nm in magnitude. Because the numerous unfolding events in the mutant protein occurred at a Ca^2+^ level likely to saturate Ca^2+^ binding sites in the inter-domain linker regions, they likely stemmed from instability in EC5, where the mutated residue was located. The frequency of total unfolding length per cycle peaked at 8.8 nm, 50.0 nm, and 66.3 nm (Figure 3C). The 50.0 nm unfolding events exceeded the size expected for the unfolding of an entire EC domain or the PICA domain, 33-45 nm, but are smaller than expected for the unfolding of two domains. This suggests that some intermediate unfolding event gave rise to this class. The 66.3 nm class of events might reflect the unfolding of two domains, which would yield an end-to-end distance increase of 66-90 nm. Many unfolding events occurred around 30 pN (Figure 3D), suggesting that V507D has a diminished ability to resist applied force compared to the wild-type protein, which has a more uniform distribution of unfolding forces.

At a physiological Ca^2+^ concentration of 20 μM, PCDH15 V507D again underwent unfolding events of varying magnitudes (Figure 3E,F). This behavior contrasted with that of the native protein at the same physiological Ca^2+^ concentration, for which very few, small unfolding events were seen. The frequency distribution of total unfolding length within each cycle peaked at 14.2 nm, 27.3 nm, and 39.4 nm (Figure 3G). As the location of the mutation, EC5 is the domain of V507D most likely to unfold. If EC5 and its neighboring linker regions were to unfold, we would expect an increase in end-to-end distance of 39.1 nm. The class at 39.4 nm therefore might well correspond to the unfolding of EC5. It is unclear what gave rise to the smaller classes of unfolding events, though unfolding of the linker regions, which would result in a total 16 nm increase in end-to-end distance, could contribute. Although we predicted that V507D would be more unstable and exhibit more unfolding at this level of Ca^2+^, the unfolding lengths did not reach such high magnitudes as at a saturating Ca^2+^ concentration. This result suggests that V507D was extended more than the wild-type protein prior to the application of force (Figure S13D). The partial unfolding at baseline means that the unfolding we observed after applying force occurs in addition to the baseline unfolding, which explains the unexpected difference in unfolding range between physiological and saturating levels of Ca^2+^ (Figure 3H).

Like the wild-type dimer, V507D underwent numerous unfolding events in the absence of Ca^2+^ as evidenced by the illustrative cycle and multiple bright branches on the illustrative heat map (Figure 3I,J). However, the bright branch closest to the origin—which reflects the extension of the protein in the absence of any unfolding events—extended to approximately 100 nm at the highest applied forces, whereas for the wild-type protein this branch reached only 50 nm. In addition, the largest end-to-end distances achieved for V507D exceeded 200 nm, a distance greater than the 125 nm characteristic of the wild-type molecule. In many individual cycles we did not observe drastic unfolding events (Figure 3I). Instead, the molecule appeared quite extended at baseline and lengthened easily with the application of force. This result suggests that the protein was already unfolded, or misfolded, at this concentration of Ca^2+^ even in the absence of force.

In the absence of Ca^2+^, although sometimes as much as 150 nm unfolded in a single cycle for V507D, the total unfolding length varied widely and the frequency peaked at 49.3 nm (Figure 3K). The average total unfolding per cycle suggests that at least one full domain unfolded during the extension phase of a cycle. At this concentration, we were likely observing unfolding events similar to those at the higher levels of Ca^2+^, along with additional events stemming from the complete lack of Ca^2+^ binding at the linker regions. There was again no clear relationship between size of unfolding events and the force at which they occurred, but many unfolding events of different sizes occurred around 30 pN (Figure 3L). Although by these metrics the behavior of V507D in the absence of Ca^2+^ resembled that at higher concentrations of Ca^2+^, the end-to-end distance range of the protein exceeded that at other levels of Ca^2+^. This result suggests that, in the absence of Ca^2+^, V507D did not fold properly even at low resting forces but was already extended at the resting force of 1 pN. Because V507D displayed behavior that was not entirely captured by our modeling approach, we next sought a method of classifying the extension patterns across different experimental conditions.

### Clustering of data by conformational state

To understand PCDH15 in various concentrations of Ca^2+^ with or without the V507D mutation, we analyzed all results from each construct and condition. The data fell into six classes, which we term “states” because, we believe, they represent different conformations of PCDH15 (Figure 4A). The defining characteristics of each state are the average value of its *x_E_* parameter and the difference between that value and those of the contiguous states (Figure 4B). Notwithstanding the substantial overlap in *x_E_* values of trajectories that we classify as belonging to different states, the average *x_E_* values clearly differ. For example, the difference between the average *x_E_* values for the first two states is 12.5 ± 0.3 nm (mean ± SEM, *n =* 2314 trajectories for state 1; *n =* 1826 trajectories for state 2), a distance consistent with the range of unfolding expected from the inter-domain linker regions. Because the difference between states 1 and 2 is smaller than what we would expect for unfolding of an entire EC cadherin, we call states 1 and 2 the folded states—that is, the states in which all EC domains and the PICA domain apparently remain folded. The differences between states 2 and 3 and between states 3 and 4 are respectively 29.9 ± 0.7 nm (mean ± SEM, *n =* 1372 trajectories for state 3) and 30.0 ± 1.3 nm (mean ± SEM, *n =* 692 trajectories for state 4), values consistent with the expectation for unfolding of a complete EC or PICA domain. The difference between states 4 and 5 is 17.6 ± 2.0 nm (mean ± SEM, *n =* 585 trajectories for state 5), a value that does not accord with unfolding of a whole domain but might represent partial unfolding of a domain. Finally, the difference between states 5 and 6 is 72.2 ± 6.0 nm (mean ± SEM, *n =* 123 trajectories for state 6), a value similar to that expected for the unfolding of two entire EC or PICA domains.

**Figure 4.**
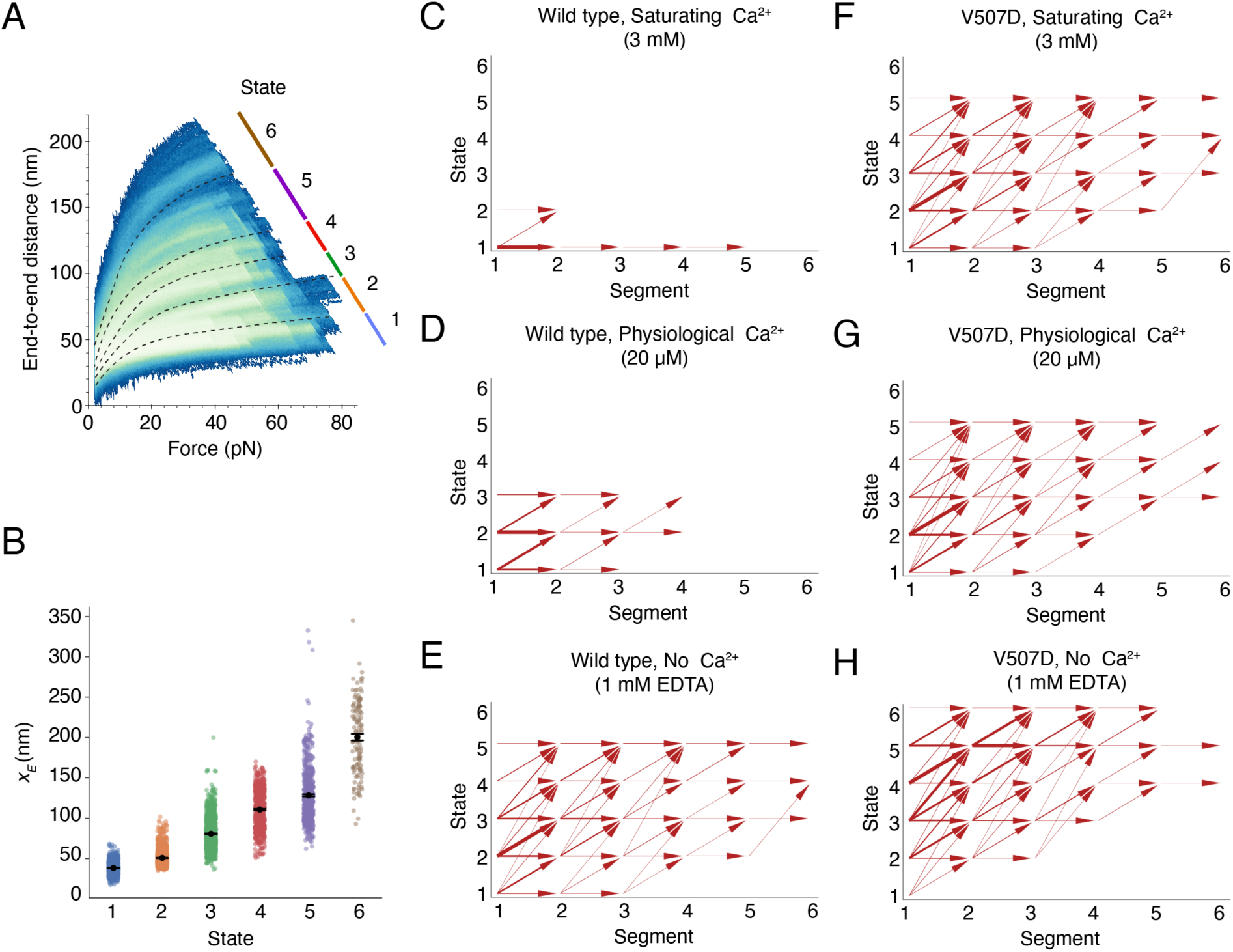
Clustering of force-ramp trajectories and inter-state transitions. (A) In a heatmap representing the relaxation-phase trajectories from all Ca^2+^ conditions and both PCDH15 constructs the lines with different colors demarcate the six conformational states. (B) The fitted *x_E_* values for individual trajectories at different conformational states are shown in the scatter plot. The data represent means ± SEMs. (C-H) Conformational transition maps summarize the trajectories of the two PCDH15 constructs at different Ca^2+^ concentrations. Segmented by conformational changes, the values along the abscissa denote individual segment within a trajectory in ascending order of force. The thickness of each arrow shaft denotes the frequency of a specific state transition. The horizontal arrows indicate conformational changes within a state, whereas the arrows pointing up and to the right denote conformational changes made between distinct states.

To understand the effect of the Ca^2+^ concentration and V507D mutation on how PCDH15 traversed the accessible state space, we segmented a molecule’s trajectory during each cycle by the conformational changes that occurred and assigned each segment to a state through *k*-nearest-neighbor classification (*k* = 3; see Supplementary Information for details). To generate transition maps for each construct and condition, we analyzed the percentage of time that the molecule spent in each state and examined the states visited by subsequent segments within one cycle. In saturating Ca^2+^ conditions, the wild-type protein remained predominantly within state 1, with a few visits to state 2 (Figures 4C, S11A, S12A). At a physiological Ca^2+^ concentration, the native dimer spent much more time in state 2 and occasionally visited state 3, a behavior suggestive of unfolding of linker regions and one EC or PICA domain (Figures 4D, S11C, S12A). In the absence of Ca^2+^, the wild-type dimer progressed up to state 5, with cycles beginning primarily in state 2 or above rather than in state 1 (Figures 4E, S11E, S12A). Under this condition, entire EC domains were likely unfolding.

In contrast to the wild-type dimer in saturating Ca^2+^ conditions, V507D at the same saturating level reached state 5 (Figures 4F, S11B, S12B). At a physiological Ca^2+^ level, V507D visited up to state 5 and exhibited transition behaviors similar to those in the saturating Ca^2+^ condition (Figures 4G, S11D, S12B). In the absence of Ca^2+^, V507D spent little time in states 1 and 2 and frequently started in higher states (Figures 4H, S11F, S12B), such as state 3, in which one EC domain was apparently already unfolded. The V507D dimer in the absence of Ca^2+^ favored higher states, unlike the wild-type protein at any Ca^2+^ concentration or V507D at greater Ca^2+^ concentrations. These results suggest that V507D had up to four unfolded EC domains in the absence of Ca^2+^. For both the native dimer and V507D, as the Ca^2+^ concentration decreased, the state space explored by the protein increased, as expected from the stabilizing effect of Ca^2+^.

### Effect of Ca^2+^ concentration on refolding

At the start of a force-ramp cycle after a large unfolding event, the protein sometimes did not return to the starting position that it had occupied at the outset of the previous cycle: the protein either did not refold at all or refolded only partially within the 2 s inter-cycle resting period. To discern any patterns in the refolding ability of PCDH15 for each construct and Ca^2+^ concentration, we compared the highest state accessed in each cycle with the state of the first segment of the subsequent cycle. If the first segment of the next cycle was in state 1 or 2, we considered this cycle to be a full refolding because the average *x_E_* values in states 1 or 2 were not consistent with the complete unfolding of any EC or PICA domain. We then analyzed the percentage of cycles with full refolding across all Ca^2+^ concentrations.

For the wild-type protein, we found that at a saturating level of Ca^2+^ PCDH15 never unfolded beyond state 2, so we obtained no refolding data for this condition. At a physiological Ca^2+^ concentration, PCDH15 refolded back to state 1 or 2 in 84.2 ± 15.8 % (mean ± SEM, *N =* 4 datasets) of the instances. In the absence of Ca^2+^, however, PCDH15 refolded back to state 1 or 2 only 62.9 ± 10.5 % (*N =* 6 datasets) of the time (Table 1).

**Table 1.** Full refolding by construct and condition. The percentage of cycles in which the dimer refolded fully are those that returned to state 1 or state 2 by the beginning of the subsequent cycle after an excursion to some higher state in the previous cycle. The wild-type PCDH15 at a saturating level of Ca^2+^, 3 mM, did not visit any state higher than state 2, so there are no data on complete refolding for that condition.

| Construct | [Ca<sup>2+</sup>] | Percentage of full refolding |
| --- | --- | --- |
| Wild type | 3 mM | — |
| | 20 μM | 84.2 ± 15.8 |
| | 0 M (1 mM EDTA) | 62.9 ± 10.5 |
| V507D | 3 mM | 82.2 ± 5.7 |
| | 20 μM | 59.3 ± 8.0 |
| | 0 M (1 mM EDTA) | 24.0 ± 16.2 |

In the V507D mutant at a saturating level of Ca^2+^, PCDH15 fully refolded on 82.2 ± 5.7 % (*N =* 24 datasets) of the occasions. At a physiological level of Ca^2+^ this value decreased to 59.3 ± 8.0 % (*N =* 16 datasets) in stark contrast with the wild-type protein, for which refolding to state 1 or 2 occurred in most instances at the same concentration of Ca^2+^. Finally, in the absence of Ca^2+^, PCDH15 V507D refolded to state 1 or state 2 in only 24.0 ± 16.2 % (*N =* 4 datasets) of the cases, which signified a severe reduction in refolding ability compared to the native protein. The differences in refolding rates between wild-type and V507D dimers suggest that the point mutation disrupts the refolding ability of PCDH15. V507D dimers unfolded more often than wild-type dimers and refolded properly less often.

### Linear stiffness of PCDH15

We calculated the enthalpic stiffness for PCDH15 and PCDH15 V507D by finding the inverse spatial derivative of each cycle in the high force regime—above 30 pN, the force at which the relationship between force and extension became essentially linear—and averaging the values across all data for each Ca^2+^ condition. In 587 determinations at physiological Ca^2+^ levels, PCDH15 had an enthalpic stiffness of 6.4 ± 0.4 mN·m^-1^. If we assume that CDH23, which is approximately 2.3 times as long as PCDH15, has mechanical properties similar to those of PCDH15, its stiffness would be about 2.8 mN·m^-1^. For both proteins in series, the stiffness *K*_TL_ of the tip link can then be calculated:

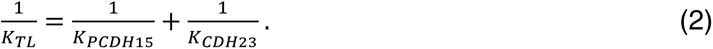

Using the values above, the enthalpic stiffness of the entire tip link in a normal animal is expected to be about 1.9 mN·m^-1^. Measurements have shown the stiffness of the gating spring to be between 0.5 mN·m^-1^ and 4 mN·m^-1^, depending on the characteristic frequency along the cochlea^15^, so our value for the enthalpic stiffness of PCDH15 lies within the expected stiffness range of the gating spring. Furthermore, the entropic stiffness of each state is less than the calculated enthalpic stiffness (Figure 5), which implies that PCDH15 is softer over the range of physiological forces than previously thought. Because the entropic stiffness of each state is lower than the calculated enthalpic stiffness, entropic elasticity is responsible for most of the mechanical response of PCDH15 over the range of physiological forces.

**Figure 5.**
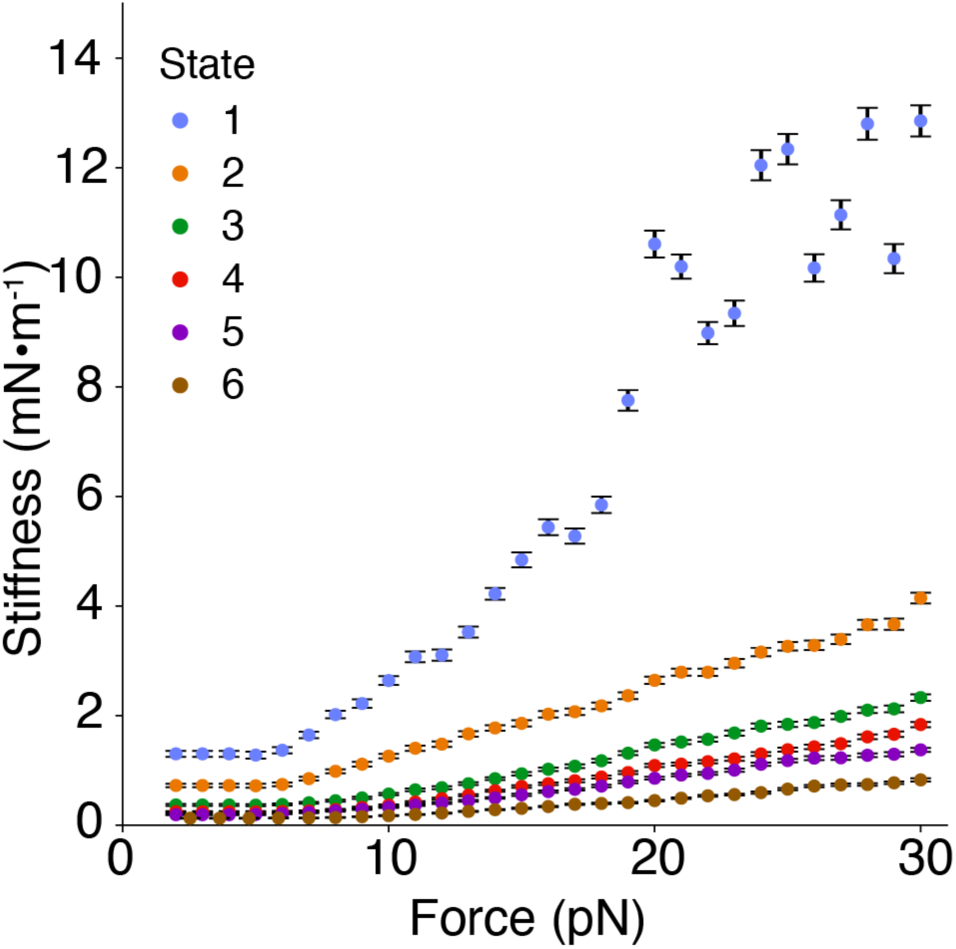
Entropic elasticity of PCDH15’s states as a function of force. The stiffnesses of the states were calculated by finding the inverse of the mean slope of each state. In state 1, when PCDH15 is fully folded, the stiffness approached the enthalpic values above 15 pN. In states 2-6, in which PCDH15 had some unfolded portions, the stiffness of PCDH15 remained below the enthalpic limit. State 1 largely comprises trajectories of the wild-type protein at a saturating level of Ca^2+^, whereas the remaining states predominantly contain trajectories at lower Ca^2+^ concentrations and from the V507D mutant. The results suggest that, under physiological conditions, the effective stiffness of PCDH15 is lower than its enthalpic limit of 40 mN·m^-1^ at a saturating concentration of Ca^2+^. Error bars represent SEMs.

## Discussion

Our findings indicate that PCDH15, in the dimeric form in which it exists within the cochlea, has the appropriate mechanical properties to serve as a portion of the gating spring. Further, at the physiological Ca^2+^ concentration and over the relevant force range, unfolding of entire EC domains is not the primary response of PCDH15 to force. Instead, the response likely comprises a series of smaller unfolding events that stem from the unfolding of the inter-domain linkers and possibly parts of EC domains.

The Ca^2+^ concentration within the endolymph ranges from approximately 20 μM at the base to 40 μM at the apex^20^. Furthermore, the local depletion of Ca^2+^ might be significant near an open transduction channel: at a distance of 7 nm, the concentration of Ca^2+^ could fall to half its maximal value. The length of a folded EC domain^30^ is about 4.5 nm, so EC10, EC11, and the PICA domain could experience a significantly lower Ca^2+^ level: the behavior of PCDH15 in the distal domains might well lie between the results for physiological levels of Ca^2+^ and those in the ion’s absence. In particular, for a Ca^2+^ concentration of 0-20 μM Ca^2+^, PCDH15 could extend around 20-100 nm. The most extreme tip link extensions observed experimentally^37^ are around 120 nm, but because mechanotransduction is very sensitive—even hair bundle deflections of 1 nm can produce a response—it is likely that the tip link ordinarily extends much less.

The V507D mutation yields significant unfolding even at a saturating concentration of Ca^2+^, and unfolds to a still more striking extent in the absence of Ca^2+^. Because the end-to-end distance at low force is far greater for the mutant dimer than for the native dimer, V507D is apparently misfolded at the baseline of 1 pN force. When the applied force is low, such as during the resting period between force-ramp cycles, the two partially unfolded strands of the dimer might tangle with each other, resulting in a misfolded protein that extends easily when force is reapplied. This behavior might occur because EC5 is more prone to unfolding due to the location of the mutation, which could then predispose the neighboring EC domains to unfold by disrupting the stability of the interdomain linkers. Furthermore, the refolding ability of V507D is significantly lower than that of the native protein at physiological and no Ca^2+^ concentrations for which the comparison is possible. If V507D is unable to refold on a timescale appropriate for normal hearing, on the order of microseconds to milliseconds, a more compliant PCDH15 dimer might underlie a softer overall tip link (Figure 6). Insufficient tension applied to the mechanotransduction channels might then be the mechanism of deafness in people with this mutation. More specifically, without an influx of Ca^2+^ through transduction channels, a hair bundle undergoes remodeling, resulting in shorter stereocilia with abnormal tip shapes^38^. Moreover, if transduction is abolished in inner hair cells, their innervation changes: they become re-innervated by inhibitory efferent neurons, which normally contact inner hair cells only before the onset of hearing^39^. If V507D cannot convey the appropriate tension to transduction channels and thus prevents their opening, changes in stereociliary structure and synaptic connectivity could underlie deafness.

**Figure 6.**
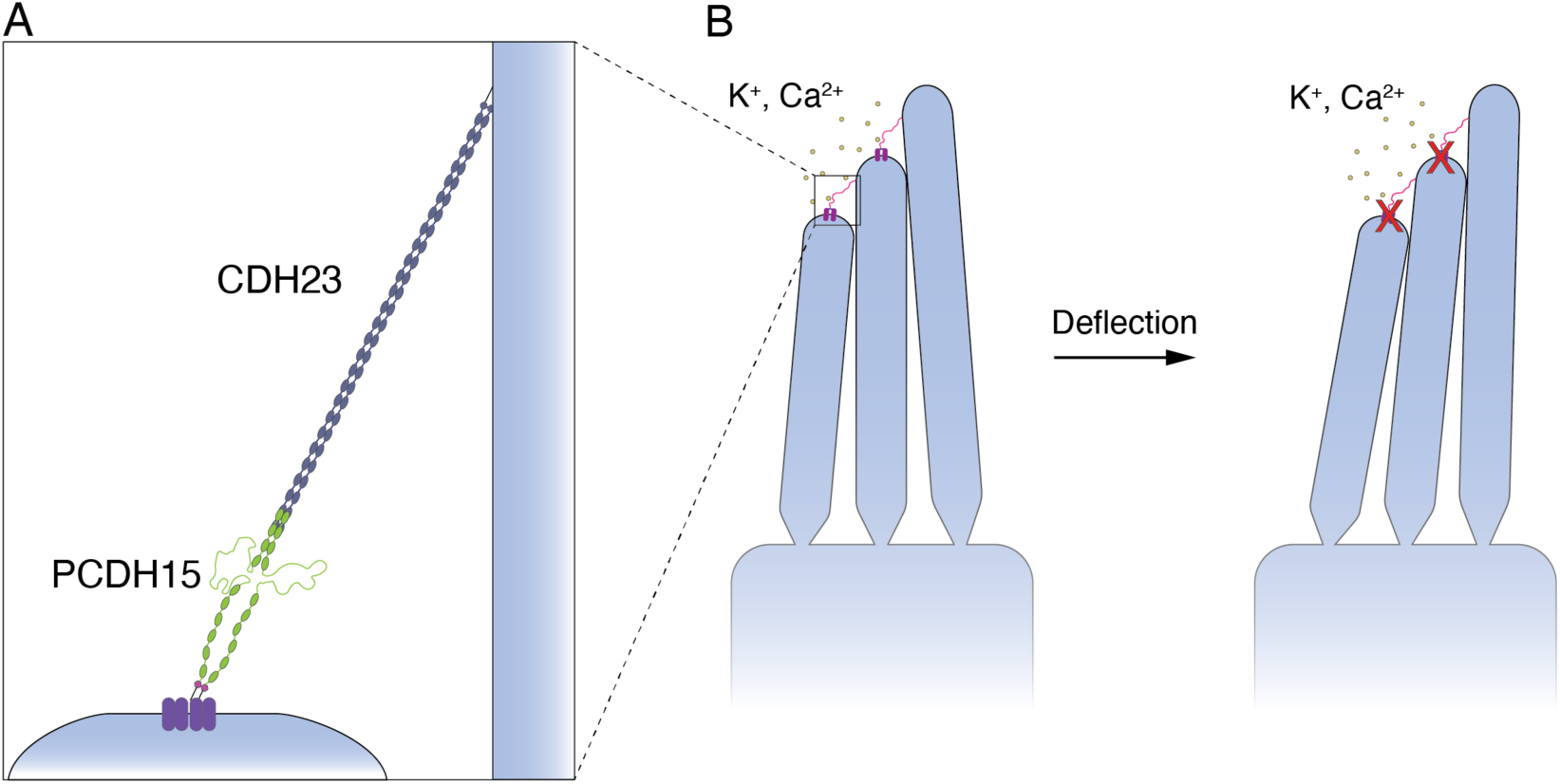
Proposed mechanism of deafness for mutated V507D. (A) Our data suggest that V507D is misfolded or partially unfolded over the physiological range of forces experienced in the cochlea. (B) If the tip links lack sufficient tension to open the mechanosensitive ion channels when stimulated, the ensuing deficit of Ca^2+^ in the stereocilia would cause their degeneration.

While the V507D point mutation disrupts the mechanics of PCDH15, individuals with this mutation retain normal equilibrium and vision, despite the expression of PCDH15 in the vestibule and eye. It is possible that the higher concentration of Ca^2+^ in the vestibular labyrinth allows mutated PCDH15 to function adequately, whereas the lower concentration of Ca^2+^ in the cochlea precludes this. It is also plausible that mutated protein can function adequately at the lower frequencies, less than 20 Hz, characteristic of the vestibular system^40^, but cannot perform well at the higher frequencies, up to 20 kHz, detected by the cochlea^2^. PCDH15 V507D might be able to refold on the timescale of low-frequency stimulation in the vestibular system, but not at the higher frequencies experienced in the cochlea. As a result, it is likely that the mutated PCDH15 cannot transmit the appropriate tension to the mechanotransduction channel, resulting in deafness in people carrying this mutation. A similar mechanism might underlie the hearing loss associated with many other mutations^16^ in PCDH15 and CDH23.

Because PCDH15 consists of repeating structural motifs, it is impossible to ascertain the specific structural origin of any unfolding event. It seems likely that, once a particular part of the molecule has unfolded, the neighboring regions become more vulnerable to unfolding, especially when Ca^2+^-binding sites are disrupted: because the Ca^2+^ ions are coordinated by residues both in the linkers and at the edges of the neighboring domains, unfolding of a neighboring domain could disrupt one or more binding sites and liberate Ca^2+^. The loss of the bound Ca^2+^ would then predispose the region to further unfolding. Although reducing the force on the protein might allow refolding to occur, proper refolding might become impossible after excessive unfolding.

These results confirm that PCDH15 has the appropriate stiffness to form a component of the gating spring and that its physical properties can be modulated by Ca^2+^. In the case of a hearing-loss mutation, PCDH15 unfolds much more frequently, is softer than the wild-type protein, and has impaired refolding ability, three features that would likely result in inappropriate tension conveyed to the transduction channels *in vivo*. The findings concerning the V507D hearing-loss mutation underscore how the tension conveyed to transduction channels is critical for normal hearing.

## Acknowledgments

The authors thank Tobias Bartsch and Ahmed Touré for assistance with establishing and calibrating the experimental apparatus; Brian Fabella, Maria Vologodskaia, Anna Kaczynska, and Sanyukta Oak for technical assistance; and Vadim Sherman for high-precision engineering of experimental chambers. CMV was supported by NIH MSTP Grant T32GM007739 and NIDCD F30 fellowship 5F30DC020104-03 and XD by Rockefeller University Institutional Funds. AJH is an Investigator of Howard Hughes Medical Institute.

## Notes

### Competing Interest Statement

The authors have declared no competing interest.

