## Supplementary material for "Nanomechanics of wild-type and mutant dimers of the tip-link protein protocadherin 15": SI

[1] Laboratory of Sensory Neuroscience, The Rockefeller University, New York, NY 10065 USA; [2] Laboratory of Populations, The Rockefeller University, New York, NY 10065 USA; [3]Earth Institute and Department of Statistics, Columbia University, New York, NY 10027 USA; [4]Department of Statistics, University of Chicago, Chicago, IL 60637 USA; [5]Howard Hughes Medical Institute, The Rockefeller University, New York, NY 10065 USA

Contents:

Supplementary Materials and Methods

Supplementary References

Supplementary Figures and Legends

### Supplementary Materials and Methods

#### Plasmid design

Constructed by Gibson assembly[1], the wild-type PCDH15 plasmid was organized as: **signal peptide-QYDDDWQYED-AviTag-IgG hinge region-GSGSGS-PCDH15 (EC1-11 and PITA)-GSGSGS-IgG hinge region-SpyTag-8xHis** (Figure S1), in which the sequence QYDDDWQYED represents the first ten amino-acid residues of PCDH15. The V507D mutation was introduced into the background of the wild type by subcloning an insert containing the mutated residue into the wild-type vector. We used isoform 1 from *Mus musculus* (UniProt entry Q99PJ1) as the PCDH15 sequence. The native signaling peptide and the first ten residues ensured proper cellular trafficking of PCDH15 for export to the membrane. The AviTag had the sequence **GLNDIFEAQKIEWHE**. The human IgG hinge region[2] had the sequence **DKTHTCPPCPAPELLGGP** and ensured that the two strands of the dimer bound each other so that force would be distributed evenly across both strands during experiments with the optical trap. We could not rule out the possibility that incorporation of the IgG hinge region altered the mechanical response of the construct in some way. The SpyTag[3] had sequence **AHIVMVDAYKPTK**.

#### Protein purification

FreeStyle 293F cells (R79007, ThermoFisher Scientific, Waltham, MA, USA) were grown at 37 °C, 150 rpm, 8 % $CO_2$ in FreeStyle 293 expression medium with 0.1 % antibiotic and antimycotic (12338018 and 15240096, ThermoFisher Scientific, Waltham, MA, USA). When the cultures reached a density of 0.8-1.0 million cells per milliliter, they were transfected by mixing Opti-MEM I reduced serum medium (31985062, ThermoFisher Scientific, Waltham, MA, USA) with PEI MAX (24765-100, Polysciences, Inc., Warrington, PA, USA) and 250 µg plasmid per 500 mL of medium. The transfected cells were then grown for seven days.

The supernatant was harvested by centrifugation and filtered through a 0.22 µm PES filter (S2GPU05RE, Millipore Sigma, Burlington, MA, USA). Protein was purified from the supernatant by a two-step protocol. First, the supernatant was mixed with Ni Sepharose 6 Fast Flow resin (17531801, Cytiva, Marlborough, MA, USA) and incubated at room temperature for 30 min. The beads were then loaded onto a chromatography column (7372522, Bio-Rad, Hercules, CA, USA) and washed with ten column volumes of wash buffer solution (150 mM NaCl, 10 mM Tris-HCl, 20 mM imidazole, and 3 mM $CaCl_2$ at pH 7.4). The protein was eluted from the resin with elution buffer solution (150 mM NaCl, 10 mM Tris-HCl, 250 mM imidazole, and 3 mM $CaCl_2$ at pH 7.4). The peak fractions were pooled, dialyzed into imidazole-free wash buffer (150 mM NaCl, 10 mM Tris-HCl, and 3 mM $CaCl_2$ at pH 7.4), and concentrated to 500 µL. The sample was run over a Superdex 200 10/300 GL size-exclusion column (28990944, Cytiva, Marlborough, MA, USA) on an ÄKTA pure 25 L (29018224, Cytiva, Marlborough, MA, USA). The peak fractions were collected, pooled, concentrated, and run on an SDS-PAGE gel to verify size. The protein was biotinylated at the AviTag (BirA500 biotin-protein ligase reaction kit, Avidity, Aurora, CO, USA) and the biotinylation was confirmed by immunoblotting. The protein was then mixed with glycerol to create a 50% v/v glycerol stock, aliquoted into single-use tubes, and flash-frozen in liquid nitrogen for storage at -80 °C for up to one year.

**Pedestal-bead functionalization**

Pedestal beads were functionalized with SpyCatcher as previously described[1]. In the present study, 1.5 µm diameter amino silica particles (ASIP-15-10, Spherotech, Lake Forest, IL, USA) were used.

To ensure that SpyCatcher was successfully attached to the beads, we tested the functionalization using a dot-blot procedure. Beads were spotted onto a nitrocellulose membrane and allowed to dry. The peptide **Biotin-AviTag-GGGSGGGS-SpyTag**, which comprises the anchors without PCDH15, was then spotted onto the beads, followed by incubation in horseradish peroxidase (HRP)-conjugated streptavidin (SA10001, ThermoFisher Scientific, Waltham, MA, USA). Binding of the biotinylated peptide to the pedestal beads indicated successful functionalization.

**Sample preparation for single-molecule experimentation**

Each experiment was performed in a rectangular channel formed by two glass coverslips mounted onto a titanium chamber. The upper surface was a 22 mm x 22 mm #1.5 glass coverslip functionalized with *n*-hydroxysuccinimide (custom order, PolyAn, Berlin, Germany) whereas the lower was a circular, 35 mm-diameter #1.5 glass coverslip (custom order, Thorlabs, Newton, NJ, USA). Before performing an experiment, we needed to covalently attach the pedestals to the upper coverslip. This was accomplished by diluting the pedestal beads in a HEPES-buffered saline solution (HEBs; 20 mM HEPES and 100 mM NaCl at pH 7.4) and pipetting the bead solution onto the functionalized coverslip, where the amines in the SpyCatcher groups covalently bound to the *n*-hydroxysuccinimide on the coverslip. The beads were allowed to react for 5 min before any unbound beads were forcefully washed off with HEBs. A thin layer of residual liquid ensured that the surface did not dry. The chamber was then sealed by securing the lower coverslip with vacuum grease. After allowing the beads to react for at least 30 min, we added a blocking buffer solution containing 10 mg/mL sulfhydryl-blocked bovine serum albumin (100-35, Lee Biosolutions, Maryland Heights, MO, USA), 150 mM KCl, 10 mM HEPES, and either 3 mM $CaCl_2$, 20 µM $CaCl_2$, or 1 mM EDTA, depending upon the experiment, to reduce non-specific binding. The chambers were blocked overnight at 4 °C and experiments were performed the following day.

The samples were incubated with protein prior to experimentation as previously described[1]. In brief, protein was diluted into a blocking buffer solution containing 10 mg/mL sulfhydryl-blocked bovine serum albumin (100-35, Lee Biosolutions, Maryland Heights, MO, USA), 150 mM KCl, 10 mM HEBs, and 3 mM $CaCl_2$, 20 µM $CaCl_2$, or 1 mM EDTA, depending upon the experiment, and flowed into the experimental chamber. The protein was allowed to incubate for 1 hr at 4 °C to allow the SpyTag on the protein to bind to SpyCatcher groups on the pedestal beads within the chamber.

Streptavidin-coated polystyrene beads 1 µm in diameter (CP01004, Bangs Laboratories Inc., Fishers, IN, USA), which we termed probe beads, were washed three times in the blocking buffer solution to remove additives from the bead storage solution. After any unbound protein had been washed from the pedestal beads with an excess of blocking buffer solution, we flowed into the chamber a solution containing probe beads and an oxygen-scavenging system. The oxygen-scavenging system[4], which comprised 18 mM D-glucose, 1000 U/mL pyranose oxidase, and 500 kU/mL catalase (P4234 and 219261 respectively, Millipore Sigma, Burlington, MA, USA), protected against the phototoxic effects of singlet oxygen and maintained a constant pH in the sample chamber. The chamber was then sealed with vacuum grease to prevent evaporation during an experiment.

#### Photonic-force microscope

The photonic-force microscope used in these studies, with sub-nanometer spatial resolution and microsecond temporal resolution, has been described in detail[1,5]. The sample chamber was mounted onto a nano-positioning stage (Nano-PDQ, Mad City Labs, Madison, WI, USA), which allowed us to move the sample in relation to the optical traps. An 852 nm force-producing laser (DL852-500, Crystalaser, Reno, NV, USA) exerted force on the probe bead and the position of the optical trap was finely controlled using a piezoelectric nano-positioning stage (Nano-3D200 and MMP micropositioning stage, Mad City Labs, Madison, WI, USA). A 1064 nm position-sensing laser (Mephisto 400 mW, Coherent, Saxonburg, PA, USA) weakly trapped the probe bead.

The x-, y-, and z-positions of the probe bead were measured by a quadrant photodiode located in the rear focal plane of the condenser lens, which captured both the unscattered light from the position-sensing laser and the light scattered by the probe bead in the optical trap. The interference of these two signals on the photodiode gave the position of the probe bead[6]. The force and position signals were sampled at 100 kHz. The diffusion of the probe bead in the position-sensing optical trap could be depicted as a three-dimensional probability distribution of its position[5,7]. To visualize the sample chamber and beads, a light-emitting diode (DC4100, Thorlabs, Newton, NJ, USA) illuminated the sample to produce an image, sampled at 5 Hz, on the camera (pco.edge 5.5, PCO, Kelheim, Germany).

#### Calibration of photonic-force microscope

The calibration of the force-producing optical trap and the position-detection system have been described in detail[1]. The position-detection system was calibrated by trapping a probe bead deep in solution with the 1064 nm position-sensing laser. The mean squared displacement of the probe bead diffusing in the trap was calculated and compared to the theoretical expectation. The sensitivity of the detector was then calculated[8].

The spring constant of the optical trap formed by the 852 nm force-producing laser was measured by increasing the intensity of the laser in discrete intervals while calculating the power spectral density of bead motion at each intensity level. Fitting the power spectrum at each level yielded a calibration curve of stiffness as a function of intensity. The intensity of the 852 nm laser was controlled by a laser modulator (Linos LM13 5W IR, Qioptiq, Goettingen, Germany).

**Force-ramp experiments**

The protocol that we used for force ramp experiments has been described[1]. We started each experiment by trapping a probe bead deep in solution with the 1064 nm position-sensing laser. We next calibrated the stiffness of the force-producing laser and the sensitivity of the position detector as described above. The probe bead was brought 250 µm from the surface of the functionalized coverslip so that the equators of the pedestal and probe beads were at the same height. We approximated the probe bead to the pedestal bead and performed an offset calculation to account for the influence of the pedestal bead on the position signal of the probe bead[5]. The probe bead was then allowed to diffuse along the edge of the pedestal bead. If a tether formed, the diffusion profile of the probe bead became severely constricted[5].

Before beginning data collection, we initiated stage drift correction by focusing the camera (pco.edge 5.5, PCO, Kelheim, Germany) on a single pedestal bead elsewhere in the field of view, but far from the tether. By tracking the movement of this distant pedestal bead, we could correct the position of the nano-positioning stage upon which the sample chamber was mounted. The optical trap of the force-producing laser was displaced by 200 nm along the axis of force application, and a brief pulse of force was delivered to the tether to break any non-specific bonds. We then initiated the force-ramp protocol, which applied force at constant rates of increase and decrease during respectively the extension and relaxation phases of each cycle (Figure S2A). The extension and relaxation phases each lasted 0.35 s. The loading rate of force application was approximately 150 $pN \cdot s^{-1}$. As force was applied, the end-to-end distance of the construct was measured (Figure S2B). Between successive cycles, the construct was held for 2 s at a low resting force of 1 pN to allow time for the protein to refold. The end-to-end distance of the construct during the low force inter-ramp intervals was approximately Gaussian-distributed (Figure S2C).

**Control of non-specific bead interactions**

For appropriate interpretation of single molecule data, it was important to ensure that the probe and pedestal beads did not adhere to one another non-specifically. We therefore tested whether the streptavidin-coated probe beads (CP01004, Bangs Laboratories, Fishers, IN, USA) would bind to the SpyCatcher-coated pedestal beads. Out of 33 attempts at tether formation over three experiments on different days, none resulted in a tether. This result largely excluded the possibility that we were measuring non-specific bead interactions.

**Characterization of the behavior of the anchors alone**

For reasons discussed previously[5], we used short and relatively stiff anchors to attach PCDH15 to the pedestal and probe beads for experimental manipulation in our optical trap. Characterizing the behavior of the anchors alone was necessary to disentangle the separate contributions of the anchors and of PCDH15 to the recorded data. We performed several control experiments on a construct including only the anchors—**Biotin-AviTag-GGGSGGGS-SpyTag**—and confirmed that it was far stiffer than the PCDH dimer (Figure S3A,B).

**Determination of the zero position of extension**

To accurately determine the extension of the protein in series with the molecular anchors, we first had to determine the zero position of extension. For each dataset, we concatenated the position signals when the construct was held at a low force of 1 pN during the inter-ramp periods. The position signals at this low resting force were approximately Gaussian in distribution, so we set the zero position three standard deviations below the mean position of the construct at the low inter-ramp resting force.

**Statistics of tether formation**

We wished to minimize the possibility that the data we recorded corresponded to multiple protein tethers. To accomplish this, we modeled the formation of tethers as a Poisson process to calculate the probability of forming only one tether during a given interval. The probability of an event happening *k* times in an interval is given by

$$P(k) = \frac{\lambda^k e^{-\lambda}}{k!}, k = 0, 1, 2, \dots . \quad \text{(S1)}$$

Here λ is the mean number of tethers in the interval, which we could not measure directly. To have 90 % probability of getting a single tether when a (single or multiple) tether is formed, we first solved for λ in the following relation:

$$\frac{P(k = 1)}{P(k > 0)} = \frac{\lambda e^{-\lambda}}{1 - e^{-\lambda}} = 0.9 . \quad \text{(S2)}$$

We obtained λ = 0.207. We then used λ = 0.207 to calculate the probability of forming a possible tether, *P(k > 0)*, regardless of number of actual tethers:

$$P(k > 0) = 1 - e^{-\lambda} . \quad \text{(S3)}$$

The result is $P(k > 0) = 0.187$. In sum, if 18.7 % of tethering attempts result in the formation of at least one tether, the probability of this tether being a single tether is 90 %. We therefore adjusted the concentration of protein empirically so that roughly one in five binding attempts resulted in a tether.

**Identifying conformational changes**

The algorithm for conformational change detection was previously described in detail[1]. Each dataset was first split into single cycles, and each cycle further split into its constituent extension and relaxation phases. The data were then smoothed using a Savitzky-Golay filter with a window of 101 points, which reduced the temporal resolution of the data from 10 µs to 1 ms. A conformational change was then detected at point *i* in the data if

$$\left|\langle x\rangle_{before} - \langle x\rangle_{after}\right| > 4\,\frac{\sigma_{before}+\sigma_{after}}{2}, \quad \text{(S4)}$$

in which $\langle x\rangle_{before}$ and $\langle x\rangle_{after}$ are the average extension values of a window of 1000 points of the smoothed data before and after point *i,* respectively, and $\sigma_{before}$ and $\sigma_{after}$ are the corresponding standard deviations. If the difference in the average extensions before and after point *i* exceeded four times the average standard deviations of the position before and after point *i*, then a conformational change was called.

**Fitting force-ramp data with a saturation model and enthalpic-stiffness term**

To quantify unfolding changes, we used a saturation model[9] with a Hookean spring term to model the data (Eq. S5, identical to Eq. (1) in the main text). In this model, the entropic extension of PCDH15 is given by the first, saturable term and the enthalpic stiffness limit is given by the second, linear term, the extension of a Hookean spring:

$$x = \frac{x_E}{1+\frac{F_{HALF}}{F}} + \frac{F}{K} \qquad \text{(S5)}$$

The maximal entropic extension of PCDH15 is given by $x_E$; $F_{HALF}$ is the force at which entropic extension is halfway complete. The end-to-end extension of the molecule is $x$. The contribution of enthalpic stiffness is given by the second term, the extension of a linear spring of stiffness $K$ under force $F$. The enthalpic spring constant $K$ was determined for each $Ca^{2+}$ concentration by averaging the inverse spatial derivatives at forces exceeding 30 pN for every cycle in all datasets in a particular condition.

To compute the size of the unfolding events, each force-ramp cycle was first split into extension and relaxation phases, then further segmented by the unfolding events that occurred. The first segment, before any unfolding occurred, was fitted for $x_E$ and $f_{HALF}$, with $K$ set to the average value for the condition to which the dataset belonged. For the subsequent segments within the same phase of a cycle, $f_{HALF}$ was held constant at its first segment value while fitting for $x_E$ alone. The size of the unfolding events was then estimated as the difference between the $x_E$ values of successive segments.

**Clustering and classification of conformational states**

To observe the total state space accessed by each PCDH15 dimer, we first plotted all its force-ramp relaxation phases without any conformational changes: these trajectories represent more stable states of the molecule (Figure 4A). Due to the noise in the data, we parameterized the data by using Eq. (S5) to fit all the parameters, $x_E$, $f_{HALF}$, and $K$; this procedure yielded a simpler representation of the data while retaining its essential aspects (Figure S10A). The goodness of fit was measured as a coefficient of determination ($r^2$). Using the Euclidean distance, we performed Ward-linkage hierarchical clustering on the parameterized trajectories within 1 pN to 45 pN to cluster into six classes, which we termed "states" because they reflect different molecular conformations. For each state, we calculated the average value of $x_E$. To provide visual context, we also plotted the original force-ramp trajectories of each state on the total state space (Figure S10B). Finally, using the previously clustered parameterized trajectories, we used Scikit-learn[10] to classify each conformational change-based segmented trajectory into different states by training and testing a *k*-nearest neighbor classifier with $k = 3$.

**Analysis of time spent in each clustered state**

To quantify PCDH15's preference for various conformational states, we analyzed the average time spent in each state for the wild-type protein and V507D mutant at different $Ca^{2+}$ concentrations. After classifying all the segmented force-ramp trajectories into different states, we calculated the time spent in the six states based on the sampling rate and the number of data points. The state preference in each condition was then summarized by the overall percentage of time spent in each state (Figure S12). To explore state preference as a function of force, we analyzed the time spent in each state from 2.5 pN to 37.5 pN and calculated the percentage of time spent during each successive 5 pN range (Figure S13), For example, 10 pN on the abscissa represents the time spent under forces between 7.5 pN and 12.5 pN.

**Analysis of PCDH15 refolding**

For each extension-relaxation cycle separately, we first determined the highest state accessed. Because the difference between average $x_E$ parameters in states 1 and 2 was not consistent with the unfolding of any full EC or PICA domain, transitions between those likely reflected unfolding of linker regions. If the highest state exceeded state 2, however, then we posited that an EC or PICA domain unfolded during the cycle and could potentially be refolded. For such a cycle, we then compared the highest state accessed with the state of the first segment of the next cycle. If the first segment of the next cycle was in state 1 or 2, we considered this cycle to have undergone a full refolding during the current cycle or the 2 s rest period between successive cycles. We then analyzed the percentage of cycles with full refolding for both wild-type and mutant PCDH15 constructs under all $Ca^{2+}$ concentrations.

**Calculation of the enthalpic stiffness**

Only portions of cycles before any unfolding occurred were considered in order to calculate the enthalpic stiffness of the fully folded protein. After observing that extension was linear in a high-force range, we excluded the effects of entropic stiffness by examining only data for forces of 30 pN and above. We excluded any segments corresponding to a force range less than 5 pN. We smoothed the segment by rounding all force values to two decimal places and averaging all end-to-end distance values falling within each rounded force value range. We further smoothed the segment by using a first-order Savitzky-Golay filter with a window matching the length of the segment. To find the average enthalpic stiffness values, we then calculated the inverse slope of the smoothed segment and removed outliers that were greater than or less than 1.5 times the inter-quartile range. We then averaged over all cycles in all datasets corresponding to each construct and condition. The resultant stiffness value, $k_{CONSTRUCT}$, corresponded to the entire construct—the protein in series with the molecular anchors tethering it to the beads at each end, which could be decomposed into its constituent parts using the following equation:

$$\frac{1}{k_{CONSTRUCT}} = \frac{1}{k_{ANCHORS}} + \frac{1}{k_{PCDH15}} \qquad \text{(S6)}$$

Using $k_{ANCHORS}$ equals to 4.11 ± 0.1 mN·m$^{-1}$ (mean ± SEM), which was the same for both constructs across all conditions, and solving for $k_{\mathrm{PCDH15}}$, we could then calculate the stiffness of PCDH15 alone, and we obtained values of 40.6 ± 10.3 mN·m$^{-1}$, 6.2 ± 0.4 mN·m$^{-1}$, and 5.7 ± 0.3 mN·m$^{-1}$ for wild-type PCDH15 at saturating (3 mM $Ca^{2+}$), physiological (20 µM $Ca^{2+}$), and no $Ca^{2+}$ (1 mM EDTA), respectively, and values of 14.1 ± 1.3 mN·m$^{-1}$, 4.3 ± 0.2 mN·m$^{-1}$, and 2.6 ± 0.2 mN·m$^{-1}$ for the V507D mutant at the same respective $Ca^{2+}$ concentrations (all means ± SEMs).

**Data presentation**

All data were analyzed using custom-written Python code. Scripts are available on GitHub at https://github.com/cvillasante/VDCH-2023.

**Data availability**

All data are available in a Dryad repository, doi:10.5061/dryad.stqjq2c8s.

**Supplementary figures and legends**

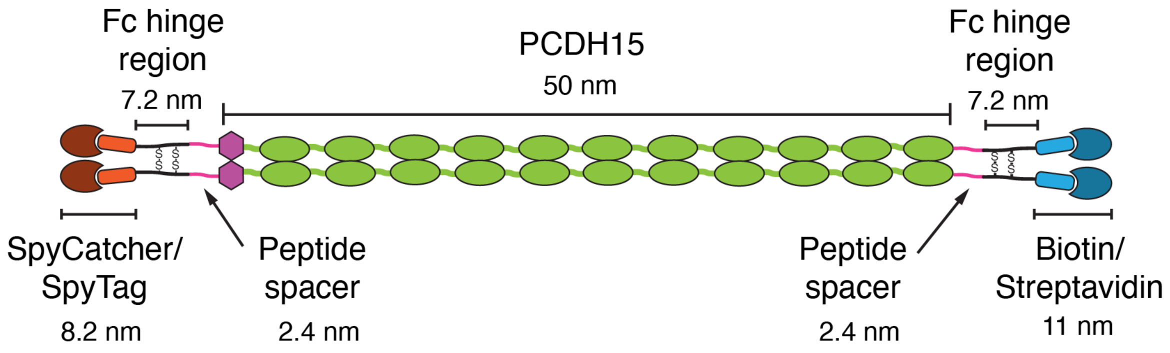


**Figure S1. Components of the construct.** In our optical trap set-up, PCDH15 was attached by means of short molecular anchors to beads on either end. The molecular anchors attaching PCDH15 to the beads on either end include a SpyCatcher[11]/SpyTag complex, one Fc hinge region from human IgG, and one GSGSGS peptide spacer on either end of PCDH15, and a biotin-streptavidin complex. The extended lengths of the non-PCDH15 components total about 38.4 nm.

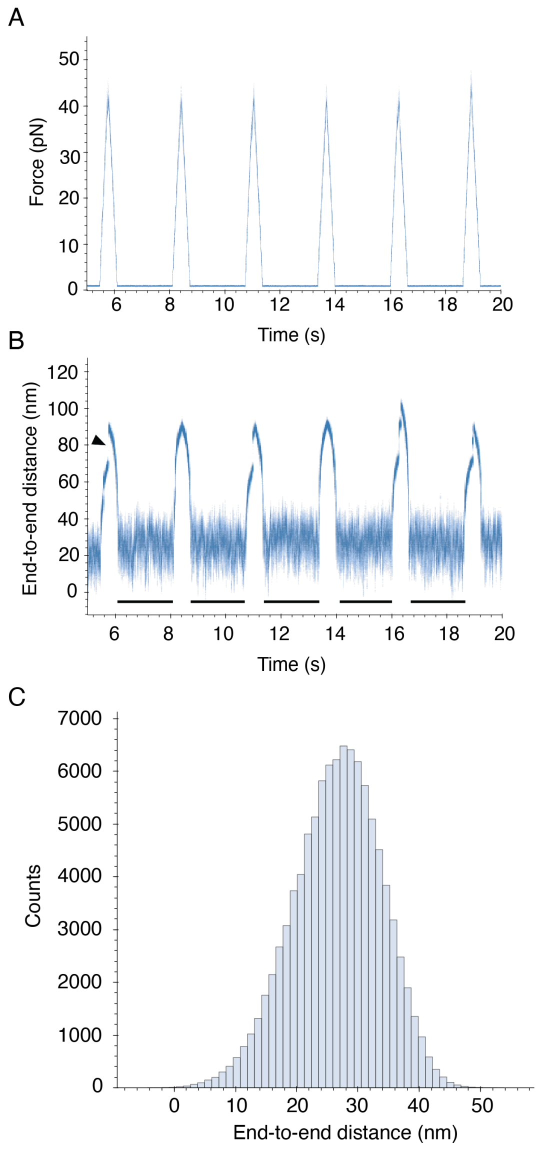


**Figure S2. Illustrative force stimulus and position traces.** (A) During a force-ramp cycle, force was first applied at a constantly increasing level up to a given maximum by increasing the spring constant of the force-producing laser. Force was then decreased at the same rate until a minimum force of 1 pN was reached. There followed a 2 s resting period during which the system is held at 1pN. (B) The corresponding position traces show the end-to-end distance of the protein in response to the force stimulus. Occasional unfolding events during the force application portions of the cycle are seen as sudden changes in the end-to-end distance; the black arrowhead marks one example. (C) The regions from panel B underlined in black correspond to the low-force inter-ramp intervals. The end-to-end distance of PCDH15 at low force was approximately Gaussian-distributed. In this case, the average end-to-end distance during the five force-free regions was 26.6 ± 0.01 nm (mean ± SEM; $n$ = 999901 data points).

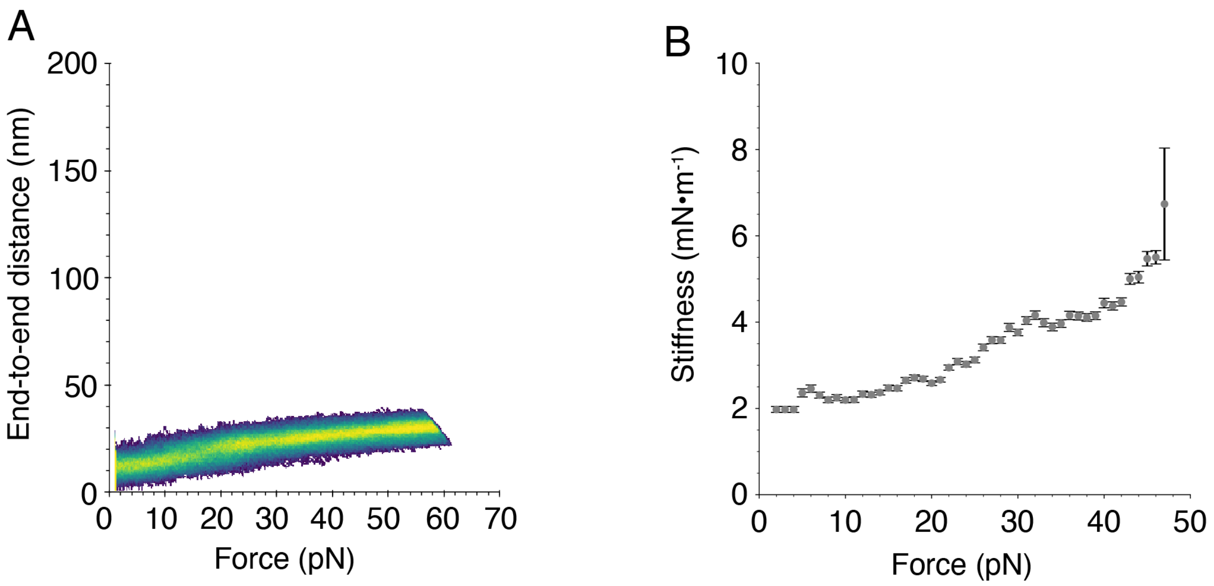


**Figure S3. Behavior of the molecular anchors in the absence of PCDH15.** (A) A representative heatmap for the anchors alone shows that they exist in one conformational state. This result confirms that the anchors do not unfold within our experimental force range. (B) We determined the stiffness of the anchors as a function of force by finding the average inverse slope of the highly occupied state space of each dataset of the anchors alone (means ± SEMs; $N = 3$).

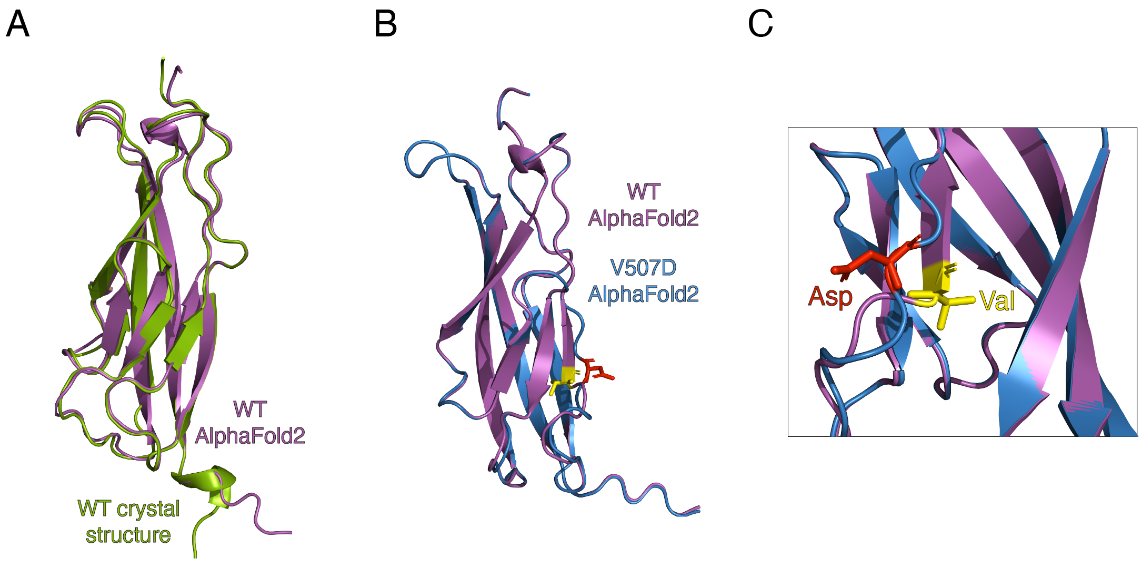


**Figure S4. AlphaFold2 prediction of EC5 with the V507D mutation.** (A) The AlphaFold2-predicted wild-type EC5 structure (purple) is shown aligned with the crystal structure of the wild-type EC5 (PDB ID: 5W1D; green). The two structures align closely, having an RMSD of 0.513 Å. (B) We then compared the structure of the AlphaFold2-predicted structure of EC5 with the V507D mutation (blue; D507 shown in red) to the AlphaFold2-predicted structure of the wild-type EC5 (purple; V507 shown in yellow). The structures are very similar, with an RMSD of 0.116 Å, but in the case of the V507D mutant, the β sheet containing the mutation site has disappeared. Because these structures have identical sequences save for the V507D mutation, this change is presumably a result of the mutation. (C) A close-up of *B* shows the wild-type V507 residue (yellow) and the mutated D507 residue (red), as well as the loss of the B strand β sheet structure with the introduction of the mutation.

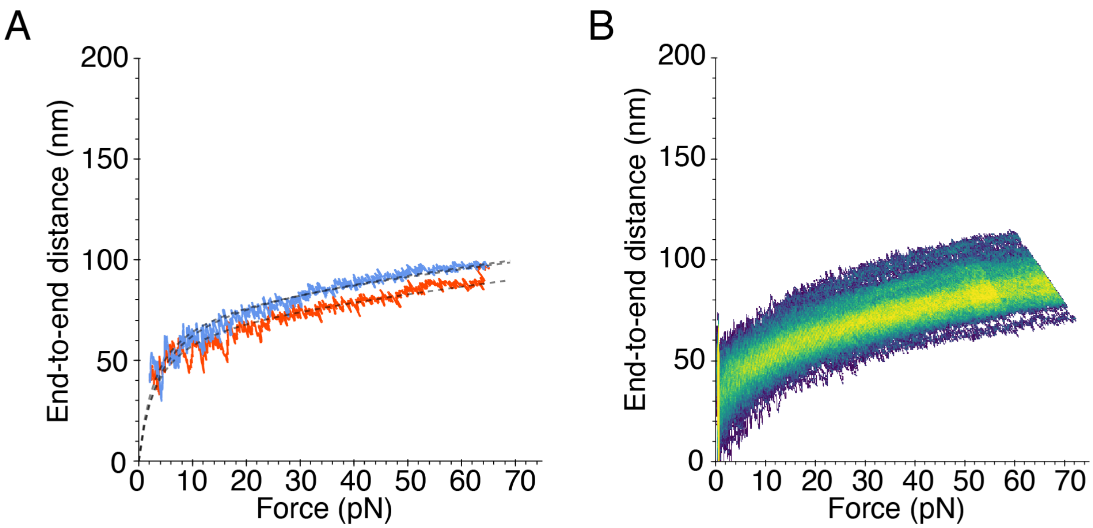


**Figure S5. A subset of V507D constructs at 3 mM $Ca^{2+}$ with only small unfolding.** (A) A subset of V507D molecules at a saturating level of $Ca^{2+}$, 3 mM, underwent only small unfolding events that could be observed at the level of individual cycles. Dashed lines represent the fit of our model, Eq. (S5), to each segment of the extension (red) and relaxation (blue) phases of a representative cycle. (B) The single bright branch on the heatmap reflects the minimal unfolding.

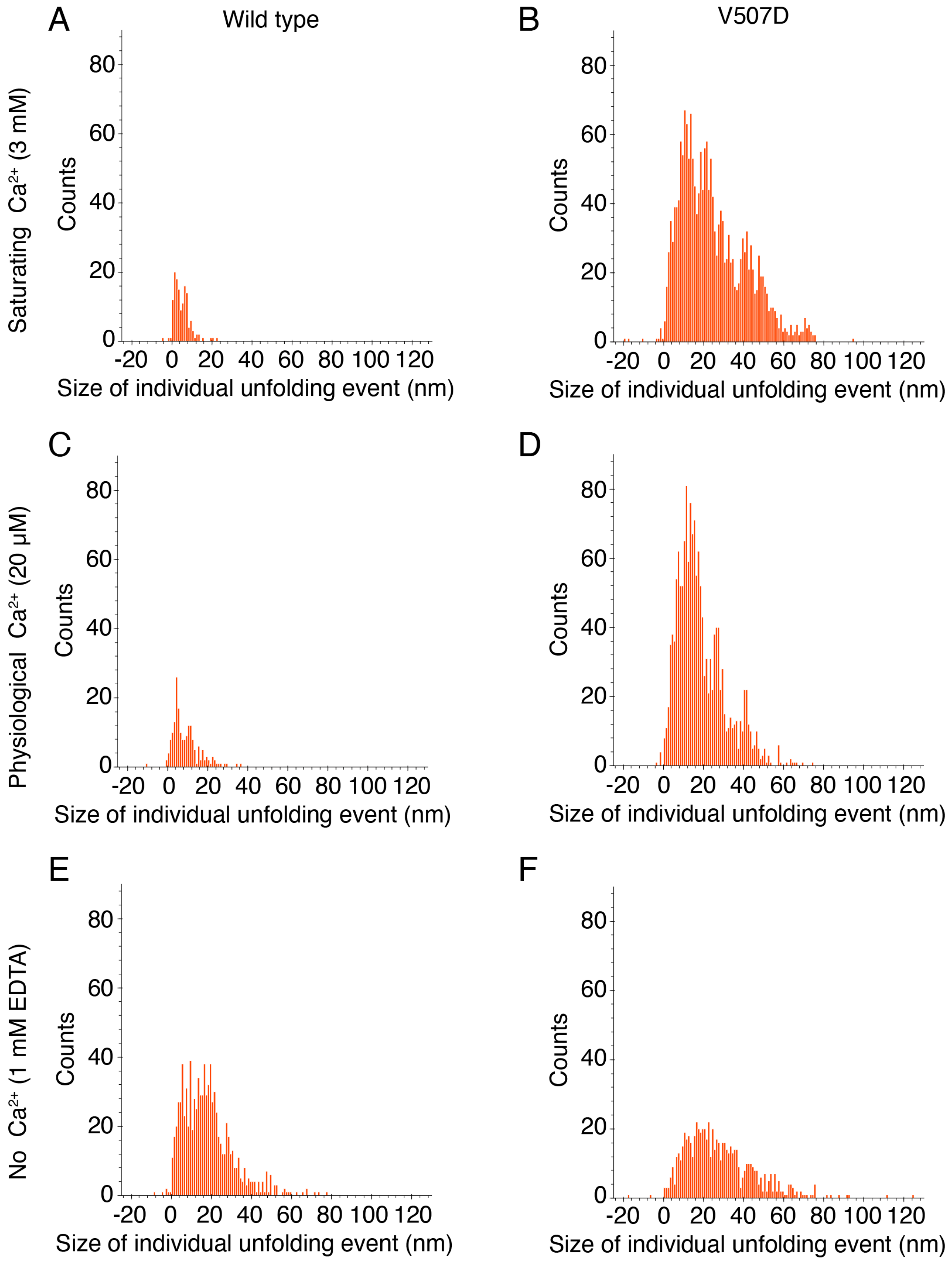


**Figure S6. Individual unfolding events for each construct and condition during the extension phase.** (A) In the wild-type PCDH15 at a saturating concentration of $Ca^{2+}$, the frequency distribution of the size of unfolding events was bimodal with peaks at 2.0 ± 0.1 nm and 6.6 ± 0.1 nm (all means ± SEMs; $N$ = 5 datasets; $n$ = 140 events). (B) In the same saturating concentration of $Ca^{2+}$, the frequency distribution of the size of individual unfolding events of

V507D constructs had many more peaks, and at higher magnitudes, than the wild type: 11.0 ± 0.1 nm, 22.4 ± 0.1 nm, 28.7 ± 0.03 nm, 32.7 ± 0.04 nm, 40.5 ± 0.1 nm, 49.5 ± 0.2 nm, and 70.5 ± 0.1 nm (all means ± SEMs; $N = 24$ datasets; $n = 1889$ events). (C) In the wild type at a physiological level of $Ca^{2+}$, the frequency distribution of the size of individual unfolding events was bimodal with peaks at 4.6 ± 0.1 nm and 8.4 ± 0.4 nm (means ± SEMs; $N = 4$ datasets; $n = 187$ events). (D) In V507D at the same physiological level of $Ca^{2+}$, the frequency distribution of the size of individual unfolding events peaked at greater magnitudes than the wild type: 12.7 ± 0.2 nm, 27.0 ± 0.1 nm, and 39.0 ± 0.2 nm (means ± SEMs; $N = 16$ datasets; $n = 1536$ events). (E) In the absence of $Ca^{2+}$, the frequency distribution of the size of the wild-type protein exhibited individual events predominantly at 4.9 ± 0.1 nm, 16.1 ± 0.2 nm, and 26.5 ± 0.4 nm (means ± SEMs; $N = 6$ datasets; $n = 868$ events). (F) In the absence of $Ca^{2+}$, the mutant V507D constructs had a unimodal frequency distribution of the size of individual unfolding events with one peak at 23.2 ± 0.5 nm (mean ± SEM; $N = 4$ datasets; $n = 694$ events).

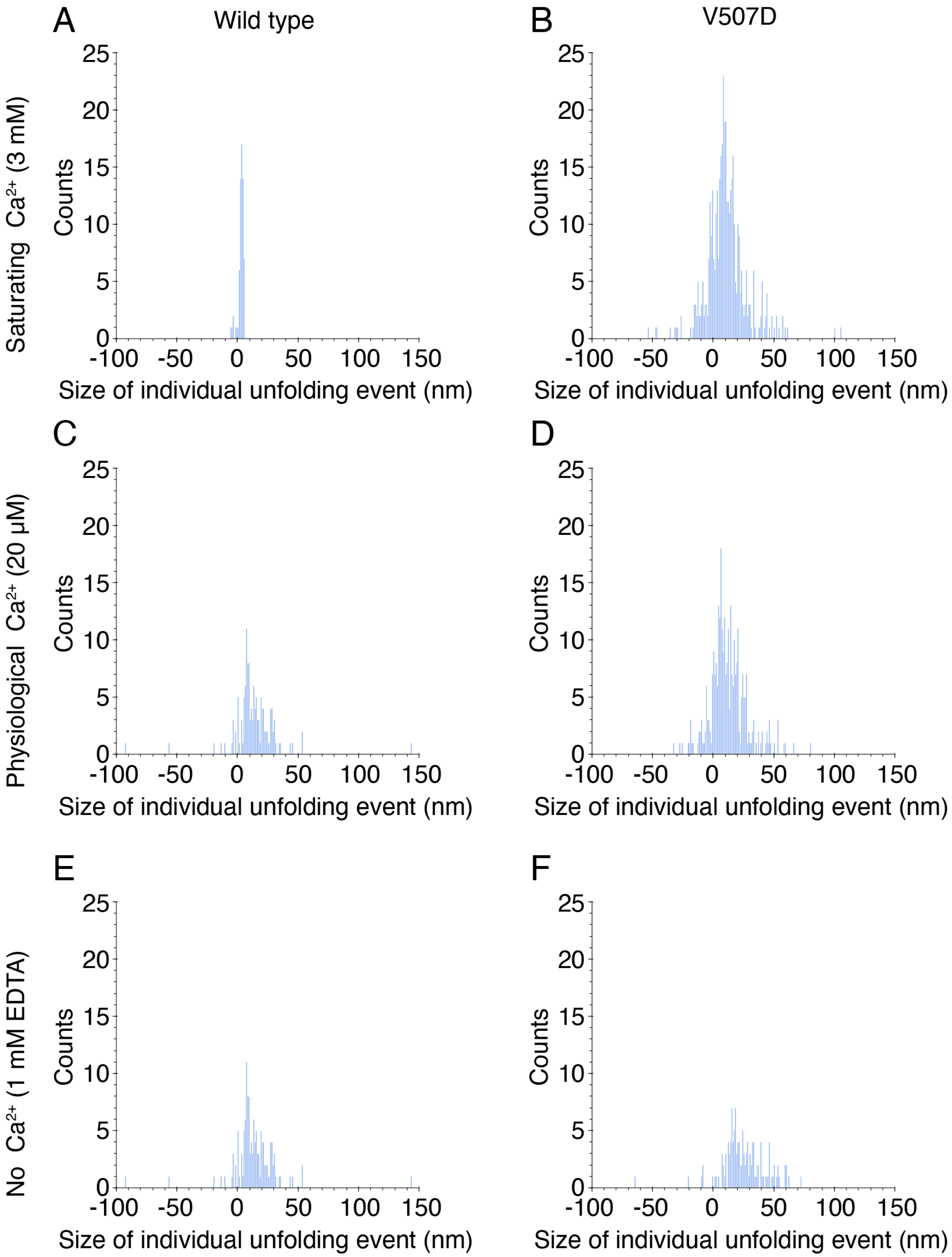


**Figure S7. Individual unfolding events during the relaxation phase.** Compared to the extension phases, fewer individual unfolding events occurred during the relaxation phases. (A) In the wild-type PCDH15 at a saturating level of $Ca^{2+}$, a peak of individual unfolding events occurred at 3.5 ± 0.2 nm (mean ± SEM; *N* = 5 datasets; *n* = 65 events). (B) At the same saturating level of $Ca^{2+}$, the V507D construct largely underwent unfolding events around 9.3 ± 0.5 nm (mean ± SEM; *N* = 24 datasets; *n* = 426 events) but with much wider variation than the wild type. (C) At a physiological level of $Ca^{2+}$, 20 µM, the wild-type protein unfolded with wide variation around a peak at 2.0 ± 1.0 nm (mean ± SEM; $N = 4$ datasets; $n = 44$ events), (D) while V507D at the same concentration mostly unfolded around 10.3 ± 0.6 nm (mean ± SEM; $N = 16$ datasets; $n = 301$ events). (E) In the absence of $Ca^{2+}$, the wild type had a peak of unfolding events at 12.0 ± 0.8 nm (mean ± SEM; $N = 6$ datasets; $n = 133$ events). (F) In V507D, also in the absence of $Ca^{2+}$, discrete unfolding events peaked at 22.2 ± 1.1 nm (mean ± SEM; $N = 4$ datasets; $n = 131$ events).

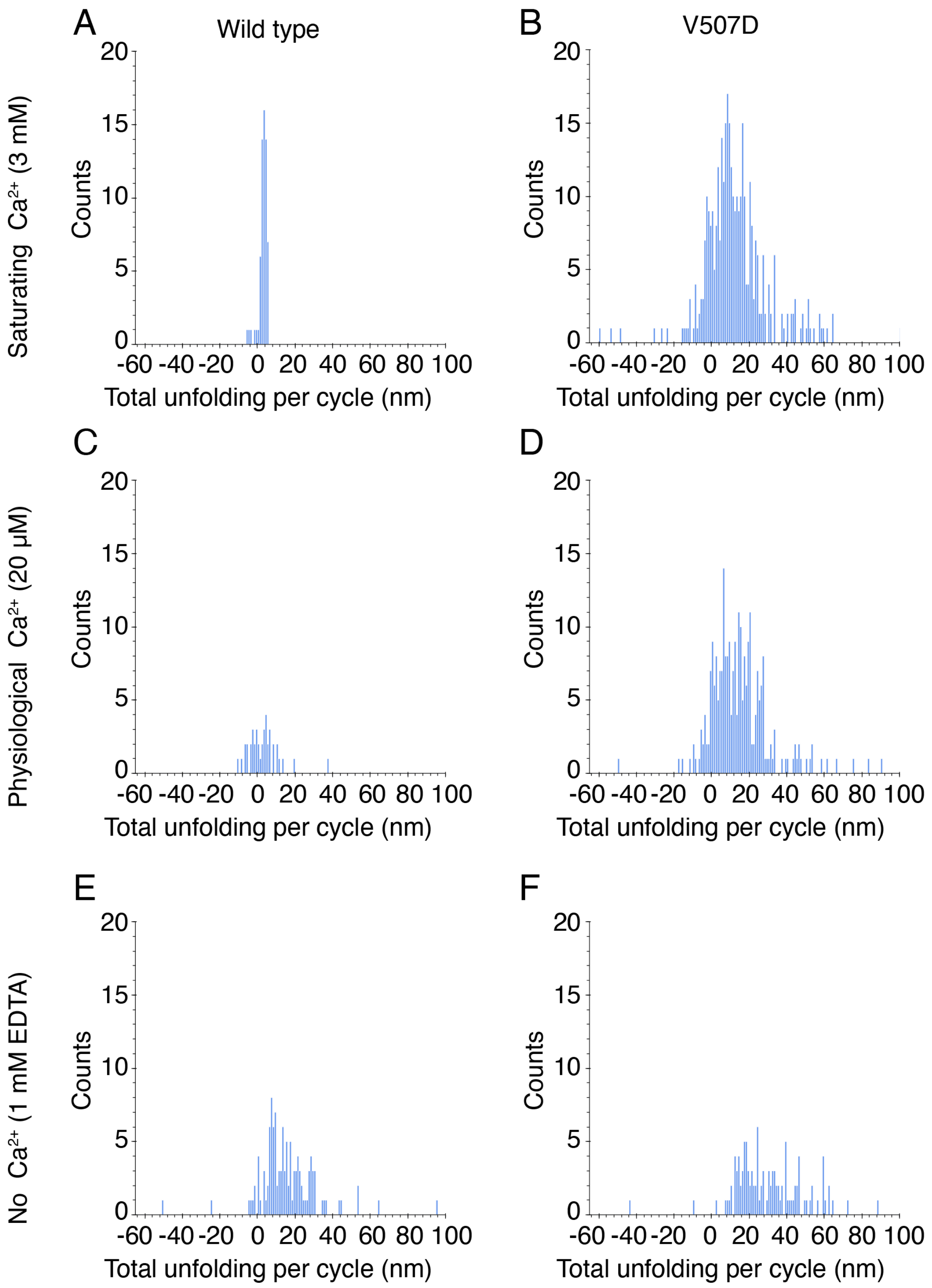


**Figure S8. Total length of unfolding per cycle during the relaxation phases** was less than during the extension phases. (A) In the wild-type protein at a saturating level of $Ca^{2+}$, the frequency distribution of the total length of unfolding per cycle during the relaxation phases was unimodal with a peak at 3.5 ± 0.2 nm (mean ± SEM; $N = 5$ datasets; $n = 63$ events). (B) In the same saturating concentration of $Ca^{2+}$, the frequency distribution of the total length of unfolding per cycle during the relaxation phases was unimodal with a peak at 9.7 ± 0.6 nm (mean ± SEM;

$N = 24$ datasets; $n = 357$ events) and had much wider variance. (C) When $Ca^{2+}$ was at a physiological concentration, in the wild type, the frequency distribution of the total length of unfolding per cycle during the relaxation phases was unimodal with a peak at 1.9 ± 0.1 nm (mean ± SEM; $N = 4$ datasets; $n = 41$ events) and had a wider variation than did wild-type constructs at a saturating level of $Ca^{2+}$. (D) At the same physiological concentration of $Ca^{2+}$ in V507D, the frequency distribution of the total length of unfolding per cycle during the relaxation phases was unimodal with a peak at 11.8 ± 0.7 nm (mean ± SEM; $N = 16$ datasets; $n = 256$ events) and a wider variance than the wild-type constructs at the same physiological concentration of $Ca^{2+}$. (E) When $Ca^{2+}$ is absent, the frequency distribution of the total length of unfolding per cycle during the relaxation phases was unimodal with a peak at 13.1 ± 0.9 nm (mean ± SEM; $N = 6$ datasets; $n = 115$ events). (F) Also in the absence of $Ca^{2+}$, in the mutant V507D, the frequency distribution of the total length of unfolding per cycle during the relaxation phases was unimodal with a peak at 26.1 ± 1.4 nm (mean ± SEM; $N = 4$ datasets; $n = 110$ events).

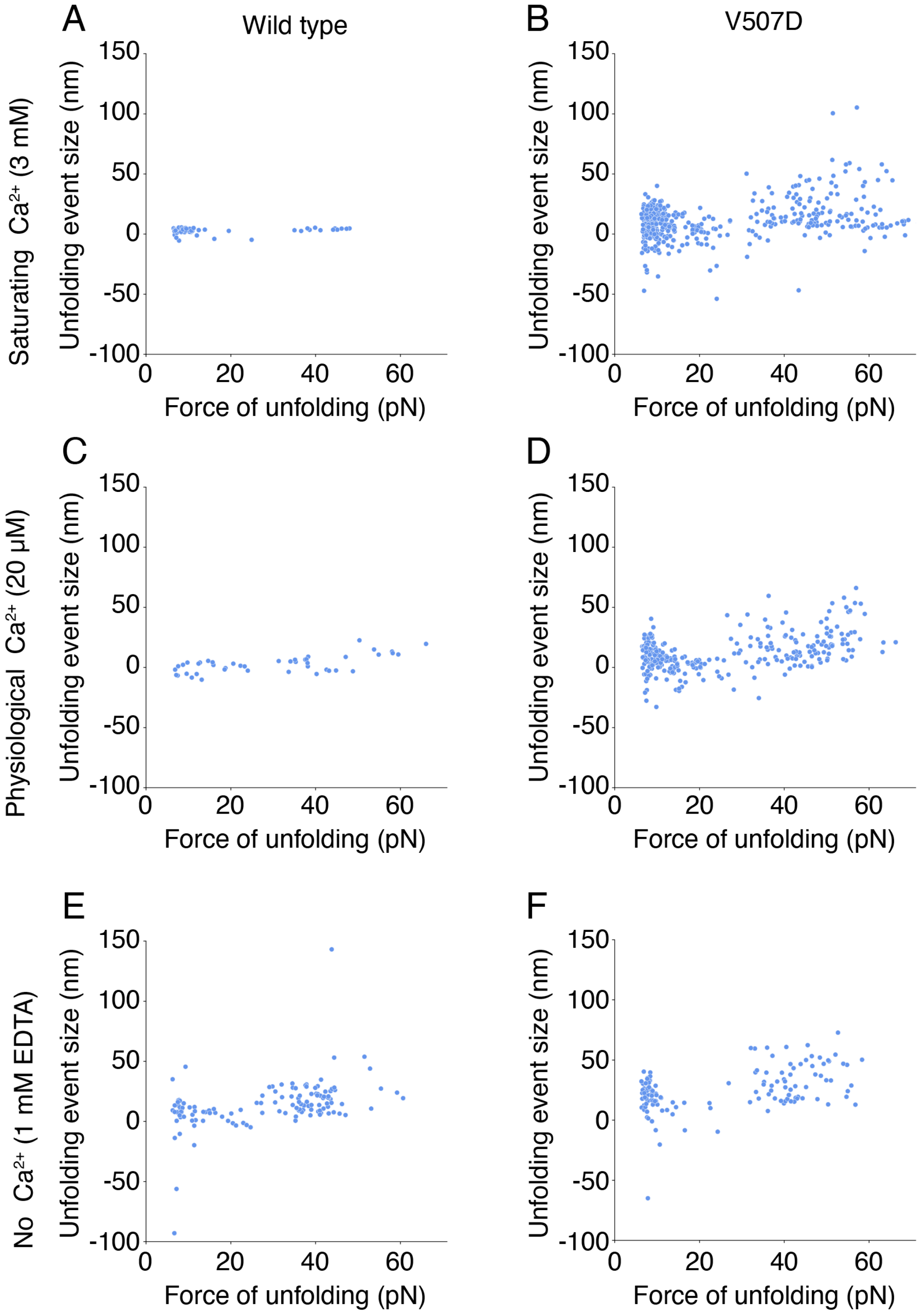


**Figure S9. The force of unfolding corresponding to individual unfolding events during the relaxation phase.** (A) In the wild type at a saturating level of $Ca^{2+}$, most unfolding events occurred at forces below 15 pN or at forces above 35 pN. (B) In the same saturating concentration of $Ca^{2+}$ in the mutant V507D construct, most unfolding and refolding events occurred at forces below 20 pN, and there was no clear association between unfolding event size and the force at which unfolding occurred. Unfolding events occurred at a relatively even distribution of forces. (C) When $Ca^{2+}$ was at a physiological level, the wild type showed no clear association between the size of the unfolding event and the force at which it occurred. (D) In V507D at a physiological level of $Ca^{2+}$, many events—both unfolding and refolding—occurred at lower forces, largely below 20 pN, again with no clear association between the size of the unfolding event and the force at which it occurred. However, compared to the wild type, relatively more unfolding events occurred at either very low or very high forces. (E) In the wild type, in the absence of $Ca^{2+}$, there was no clear relationship between the size of the unfolding event that occurred and the force of unfolding. (F) In V507D, also in the absence of $Ca^{2+}$, many unfolding events occurred below 10 pN and above 30 pN.

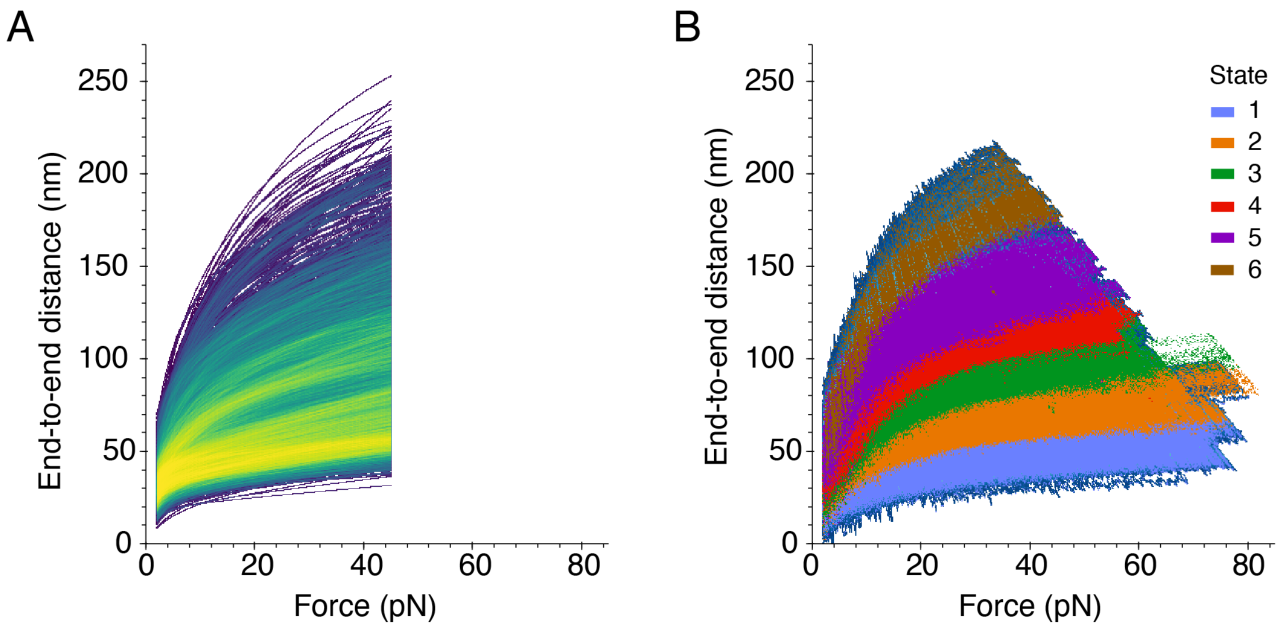


**Figure S10. Illustrative fit to the data by our model and overlay of states onto heatmap of all data.** (A) To reduce the noise in the data, we used Equation (1) or Equation (S5) to fit each relaxation trajectory from both PCDH15 constructs and all $Ca^{2+}$ conditions. The fits for all the trajectories are plotted here as a heatmap in which brighter colors represent more highly occupied regions. Fits were performed only between 1 pN and 45 pN due to the maximal forces reached by the most-extended molecules. (B) Using the fits from panel A, we performed Ward-linkage hierarchical clustering based on the Euclidean distance. The six resultant classes are shown overlaid on the heatmap of all relaxation trajectories from both PCDH15 constructs under all $Ca^{2+}$ conditions.

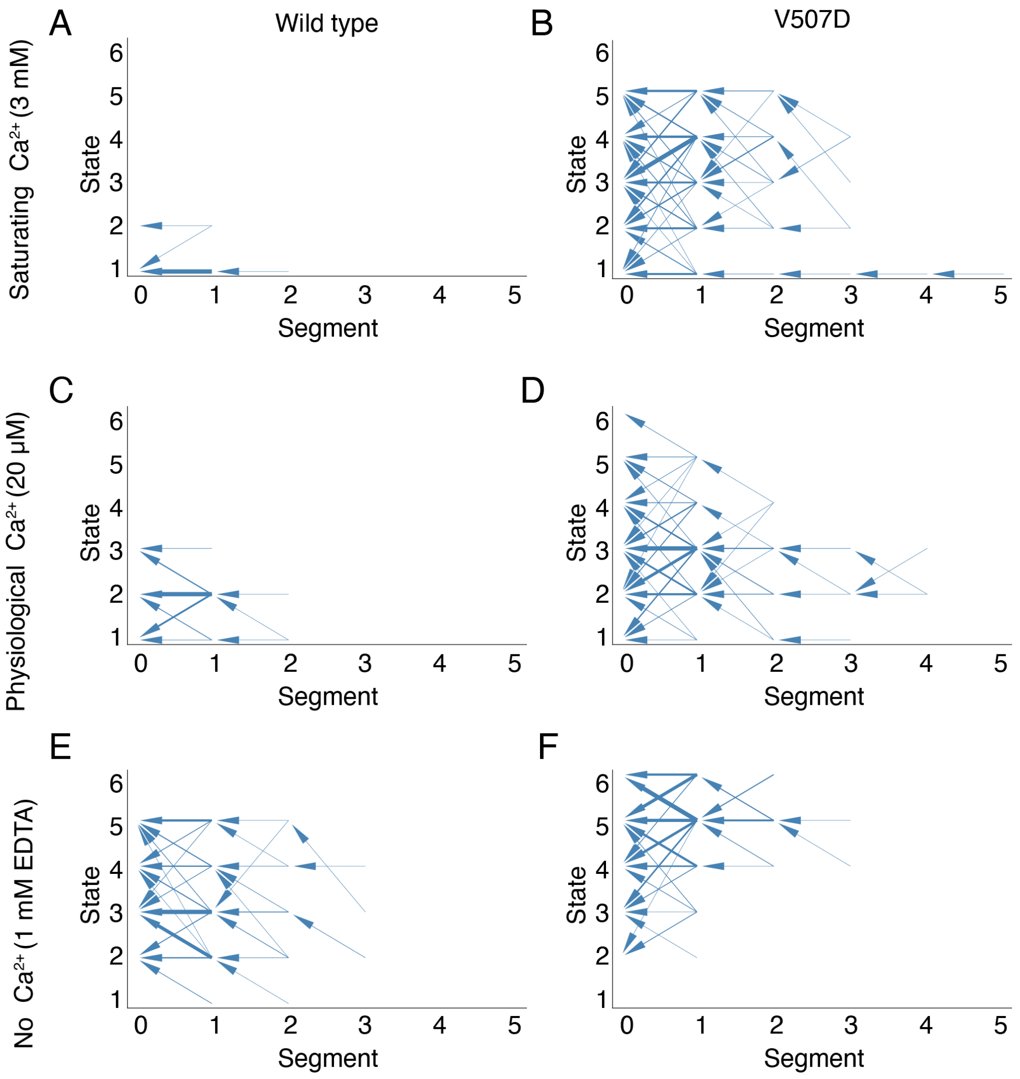


**Figure S11. Inter-state transitions during the relaxation phases.** These plots depict the inter-state transitions between consecutive segments—each segment bounded by unfolding events—during the relaxation phase of each cycle in both constructs across all three $Ca^{2+}$ concentrations. The thickness of the arrow shaft indicates the frequency of the transition. (A) At a saturating level of $Ca^{2+}$, the wild-type protein remained predominantly in state 1 during the relaxation phase. (B) In contrast, at the same level of $Ca^{2+}$, V507D existed up to state 5 during the relaxation phase. (C) At a physiological level of $Ca^{2+}$, the wild-type dimer primarily populated state 2 but explored up to state 3 during the relaxation phase. (D) In contrast, V507D at the same physiological concentration of $Ca^{2+}$ ventured up to state 6. (E) When $Ca^{2+}$ was absent, the wild-type protein remained predominantly in state 3, though it explored up to state 5 during the relaxation phase of the cycle. (F) In the absence of $Ca^{2+}$, the mutant V507D protein did not visit state 1 during the relaxation phase, but primarily inhabited states 5 and 6.

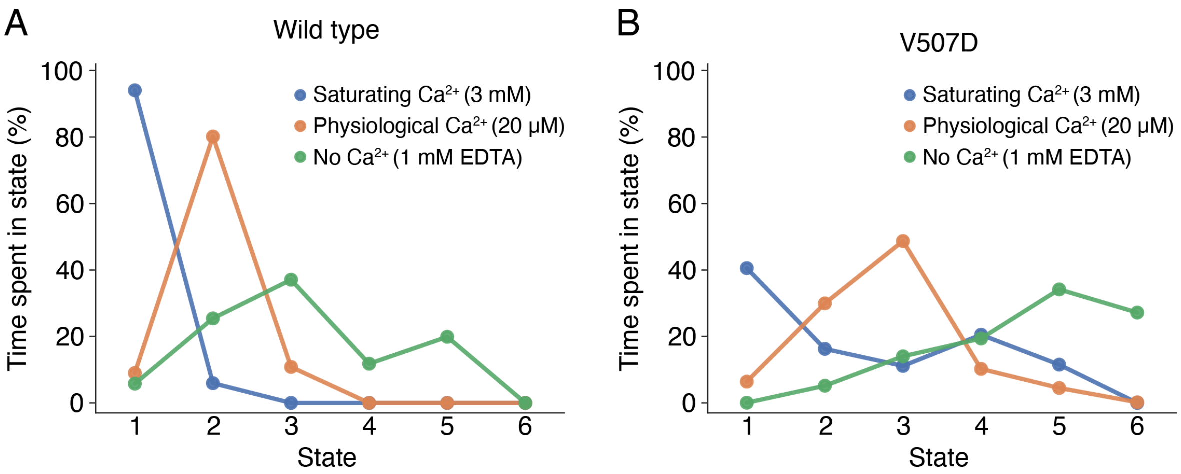


**Figure S12. Overall percentage of time spent in each state.** (A) At a saturating concentration of $Ca^{2+}$, wild-type PCDH15 remained predominantly in state 1. When $Ca^{2+}$ was at a physiological level, the wild type occupied state 2 most often. When $Ca^{2+}$ was absent, the wild type most frequently existed in state 3, but states 4 and 5 were occupied as well. (B) At a saturating concentration of $Ca^{2+}$, V507D existed primarily in state 1, but for only about 40 % of the time compared with nearly 100 % of the time for the wild-type dimer. At a physiological concentration of $Ca^{2+}$, V507D most frequently occupied state 3, unlike the native protein that most frequently existed in state 2. In the absence of $Ca^{2+}$, V507D spent the most time in states 5 and 6, whereas the wild-type protein remained most frequently in state 3.

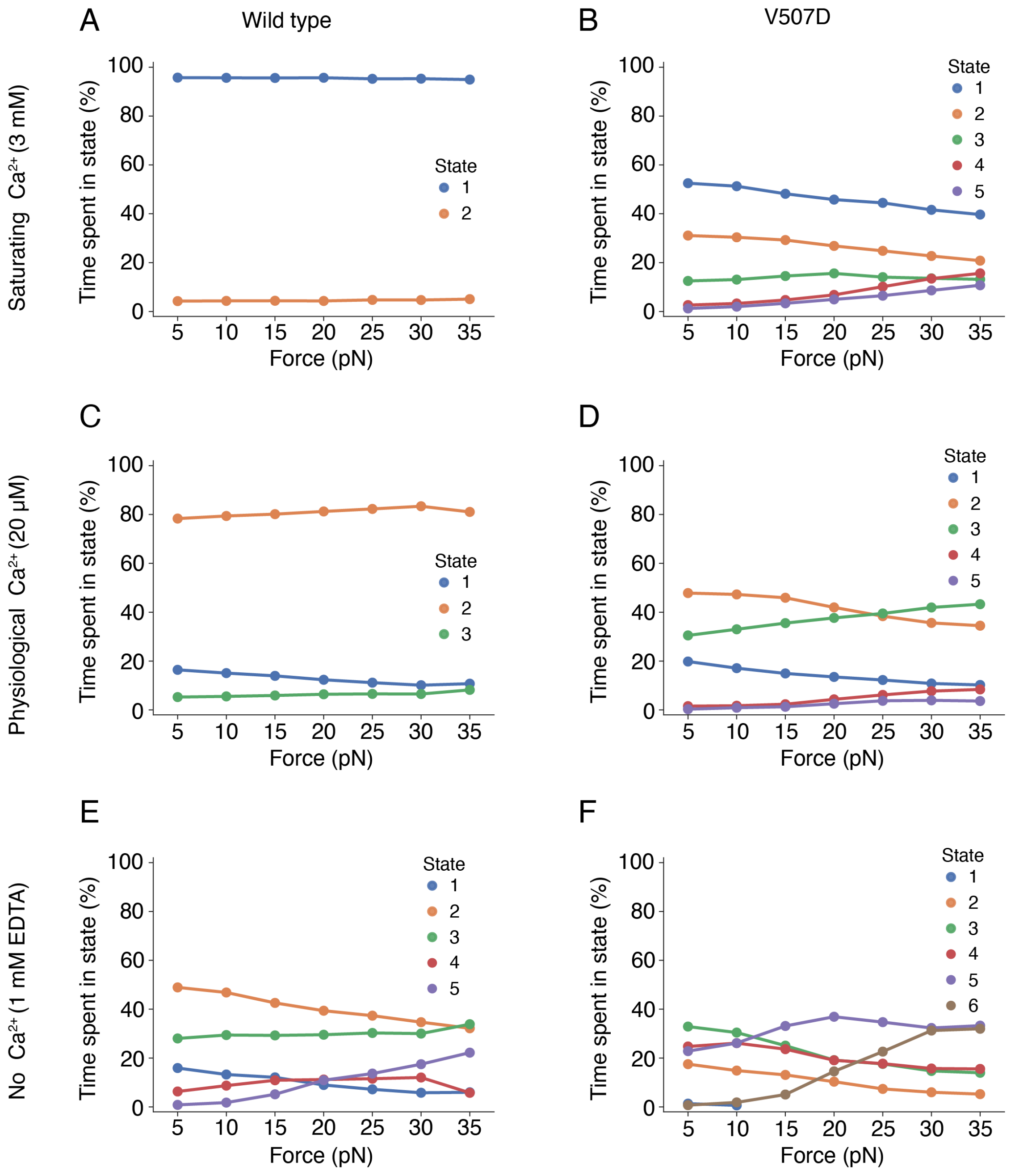


**Figure S13. Percentage of time spent in each state as a function of force during the extension phase.** (A) At a saturating concentration of $Ca^{2+}$, the wild-type PCDH15 existed nearly 100 % of the time in state 1 across the force range explored. (B) In contrast, V507D existed only about half of the time in state 1 across all forces even though it was still the most occupied state in the mutant. (C) At a physiological concentration of $Ca^{2+}$, the wild type existed most frequently in state 2 across all forces. (D) In the same physiological concentration of $Ca^{2+}$, V507D also spent the most time in state 2, but above 25 pN it began to spend more time in state 3. (E) In the absence of $Ca^{2+}$, the wild-type protein occupied state 2 most of the time, though only about 40-50 % of the time compared to about 80 % at physiological $Ca^{2+}$. (F) In contrast, in the absence of $Ca^{2+}$, V507D primarily occupied state 3 below a force of 15 pN, above which it spent the most time in state 5. Above a force of 30 pN, its frequencies in states 5 and 6 were very similar.
